# High-dimensional single-cell definition of CLL T cells identifies Galectin-9 as novel immunotherapy target

**DOI:** 10.1101/2022.12.15.519719

**Authors:** L Llaó Cid, JKL Wong, I Fernandez Botana, Y Paul, M Wierz, A Flörchinger, S Gonder, G Pagano, M Chazotte, K Bestak, C Schifflers, M Iskar, T Roider, F Czernilofsky, Bruch P-M, JP Mallm, A Cosma, DE Campton, E Gerhard-Hartmann, A Rosenwald, D Colomer, E Campo, D Schapiro, S Dietrich, P Lichter, E Moussay, J Paggetti, M Zapatka, M Seiffert

## Abstract

Failure of cancer immunotherapy is linked to T cell exhaustion. To decipher the underlying mechanisms, we explored the T cell landscape in blood, bone marrow and lymph node samples of patients with chronic lymphocytic leukemia (CLL), and spleen samples of a CLL mouse model. By single-cell RNA-sequencing, mass cytometry (CyTOF), and multiplex image analysis of tissue microarrays, we identified a disease-specific accumulation of distinct regulatory T cell subsets and T cell exhaustion stages and their trajectories in CLL lymph nodes. Integration of T cell receptor sequencing data revealed a clonal expansion of CD8^+^ precursor exhausted T cells (T_PEX_), suggesting their CLL reactivity. Interactome analyses identified the TIM3 ligand Galectin-9 as a novel immunoregulatory molecule in CLL. Blocking of Galectin-9 in CLL-bearing mice slowed down disease development and reduced the number of TIM3-expressing T cells. Galectin-9 expression correlated with shorter survival of patients with CLL, renal cell carcinoma or glioma.

**Statement of significance:** Our findings for the first time define the T cell landscape in CLL lymph nodes and reshape the current understanding of T cell exhaustion in this malignancy. They further introduce Galectin-9 as novel immune checkpoint with a high potential to overcome resistance to PD1 targeting drugs in CLL and beyond.

## Introduction

Failure of response to immune checkpoint inhibitors (ICI) is commonly seen in cancer patients with unknown causes and predictors, limiting the use of these novel immunotherapy approaches. B cell lymphomas develop in lymph nodes (LNs), the site of T cell priming and activation in infections and cancer. Even though there is a constant interaction of malignant B cells with CD4^+^ and CD8^+^ T cells in this tissue, adaptive immune control fails and response to ICI is very limited in patients with B cell lymphomas. In particular, very low response rates to anti-PD1 antibodies were observed in patients with chronic lymphocytic leukemia (CLL) (Ding et al., 2017; Xu-Monette et al., 2018), which is a neoplasm of mature B cells that accumulate in LNs and blood. In contrast to patients with acute lymphocytic leukemia (ALL), also response rates to CD19 CAR-T cell therapy remain below the expectations in CLL (Cappell et al., 2020). A lack of fit and functional T cells is discussed as a main limitation explaining the failure of immunotherapy responses in CLL and beyond (Fraietta et al., 2018; van Bruggen et al., 2019). Chronic exposure of T cells to tumor (neo)antigens and an immune suppressive microenvironment lead to their exhaustion and dysfunction (Blank et al., 2019). Whereas terminally exhausted T cells, characterized by high PD1 expression levels and expression of TOX, fail to reactivation by ICI, their precursor state that expresses lower levels of PD1 and is positive for TCF-7, harbors self-renewal capacity and the ability to control tumor development upon ICI treatment (Im et al., 2016; Scott et al., 2019; Utzschneider et al., 2016). The presence of these cell states has been confirmed in CLL patients and mouse models (Hanna et al., 2021), even though a characterization of T cells in CLL LNs is lacking due to the limited access to respective samples. Current efforts aim at characterizing mechanisms of T cell exhaustion in cancer and at identifying novel therapeutic targets to overcome this limitation.

Here, we applied single-cell RNA sequencing (scRNA-seq) and mass cytometry (CyTOF) of LNs, peripheral blood (PB) and bone marrow (BM) samples from patients with CLL, as well as tumor-free reactive lymph nodes (rLN) from healthy controls, combined with multiplex imaging of LN tissue sections and microarrays to characterize the distribution, phenotype, and function of T cells under chronic exposure to malignant cells. We show that CLL LNs constitute a unique niche, where clonally expanded CD8^+^ T cells expressing CD39 and with exhausted phenotype are enriched in comparison to healthy LNs. We further defined cells with a signature of precursors of exhausted cells that accumulate in CLL LNs, as well as several types of regulatory T cells. The results of interactome analyses led us to target Galectin-9 in a CLL mouse model, which resulted in improved T cell function and an attenuated tumor development. Our analyses link Galectin-9 expression to survival of patients with CLL, kidney cancer or brain tumors.

## Results

### Definition of the T cell landscape in CLL using mass cytometry

To comprehensively characterize the T cell compartment associated with CLL, a large-scale and high-dimensional analysis of T cells from CLL patients as well as healthy controls (HC) was conducted using mass cytometry, scRNA-seq and multiplex imaging of tissue sections followed by integrative data analysis (Figure 1A). The patient cohort reflects the heterogeneity of CLL with respect to cell-of-origin, including both *IGHV* mutated (n=10) and unmutated (n=10) cases, as well as patients with the prevalent genetic aberrations (Suppl. Table 1). We performed mass cytometry profiling of T cells from 22 CLL LNs (including 2 patients with 2 time points), plus 7 paired PB and 3 paired BM samples, as well as 13 reactive LNs (rLNs) from HC using a panel of 42 antibodies (Key Resource Table) designed to identify and characterize naïve, memory, effector, regulatory and exhausted T cells. The analysis comprised 5.29 x10^6^ T cells, with a median of 51,937 cells per sample (Suppl. Figure 1A-C). Unsupervised graph-based clustering based on the differential expression of the analyzed proteins grouped cells into 30 clusters which are presented as uniform manifold approximation and projection (UMAP) plot (Figure 1B, C, Suppl. Figure 2A). This approach led to the identification of 15 CD4^+^ T cell clusters (including 4 T_REG_ clusters), 9 CD8^+^ T cell clusters, 4 double-negative T cell clusters, and 2 clusters with a mixture of CD4^+^ and CD8^+^ T cells (Figure 1D, Suppl. Figure 2B-C, and Suppl. Table 2). The expression of CD45RA, CD45RO, and CCR7 allowed for a general classification of T cells into naïve (T_N_, CD45RA^+^ CD45RO^-^ CCR7^+^), central memory (T_CM_, CD45RA^-^ CD45RO^+^ CCR7^+^), effector memory (T_EM_, CD45RA^-^ CD45RO^+^ CCR7^-^) and effector memory cells re-expressing CD45RA (T_EMRA_, CD45RA^+^ CD45RO^-^ CCR7^-^). The CD8^+^ T cell subsets comprised 1 naïve (CD8 T_N_), 1 central memory (CD8 T_CM_), 3 subsets with a short-lived effector cell (SLEC) phenotype expressing TBET and KLRG1 (CD8 T_EMRA_ TBET, CD8 T_EMRA_ TBET, and CD8 T_EMRA_ NCAM) (Joshi et al., 2007; Obar et al., 2011), and 4 effector memory subsets with a higher expression of the cytotoxic molecule granzyme K (GZMK) and an increasing expression of exhaustion-related molecules such as the inhibitory receptors PD1 and TIGIT, the TFs EOMES and TOX, and the ectoenzymes CD38 and CD39 (CD8 T_EM_ GZMK, CD8 T_EM_ TBET GZMK, CD8 T_EX_ CD38, and CD8 T_EX_ CD39) (McLane et al., 2019). The CD4^+^ T cell subsets comprised 1 naïve (CD4 T_N_), 3 central memory (CD4 T_CM1_ CCR7, CD4 T_CM2_ CD25, and CD4 T_CM3_), 1 follicular helper (T_FH_), and 6 effector memory subsets. Of these, two expressed TBET and KLRG1, resembling SLEC subsets (CD4 T_EM_ NCAM, and CD4 T_EM_ TBET), while 3 of them expressed higher levels of PD1 and TIGIT (CD4 T_EM_ PD1), CTLA4 and CD38 (CD4 T_EM_ CTLA4), or CD39 (CD4 T_EM_ CD39). The last CD4 T_EM_ subset expressed EOMES, GZMK and PD1 (CD4 T_EM_ GZMK), and shares therefore characteristics of T regulatory type 1 cells (T_R_1) and CD4^+^ T cells with cytotoxic properties (Oh and Fong, 2021; Roessner et al., 2021). In addition, FOXP3 expression identified 4 T_REG_ clusters within the CD4^+^ T cells, 2 with a central memory phenotype (CD4 T_REG-CM1_ and CD4 T_REG-CM2_), and 2 subsets with an activated inhibitory phenotype (CD4 T_REG_ PD1 and CD4 T_REG_ CD39). CD4 and CD8 double negative (DN) CD3^+^ T cells comprised 2 clusters with a phenotype similar to CD8 SLEC subsets (DN T_EM_ KLRG1 and DN T_EMRA_ TBET), and 2 cytotoxic clusters expressing GZMK and EOMES as well as CD38 (DN T_EMRA_ CD38) or the transcription factor HELIOS (DN T_EM_ HELIOS). A KI67^+^ proliferative subset (T_PR_), as well as a cluster with high expression of ICOS (T_EM_ ICOS) were both composed of CD4^+^ and CD8^+^ T cells.

**Figure 1:**
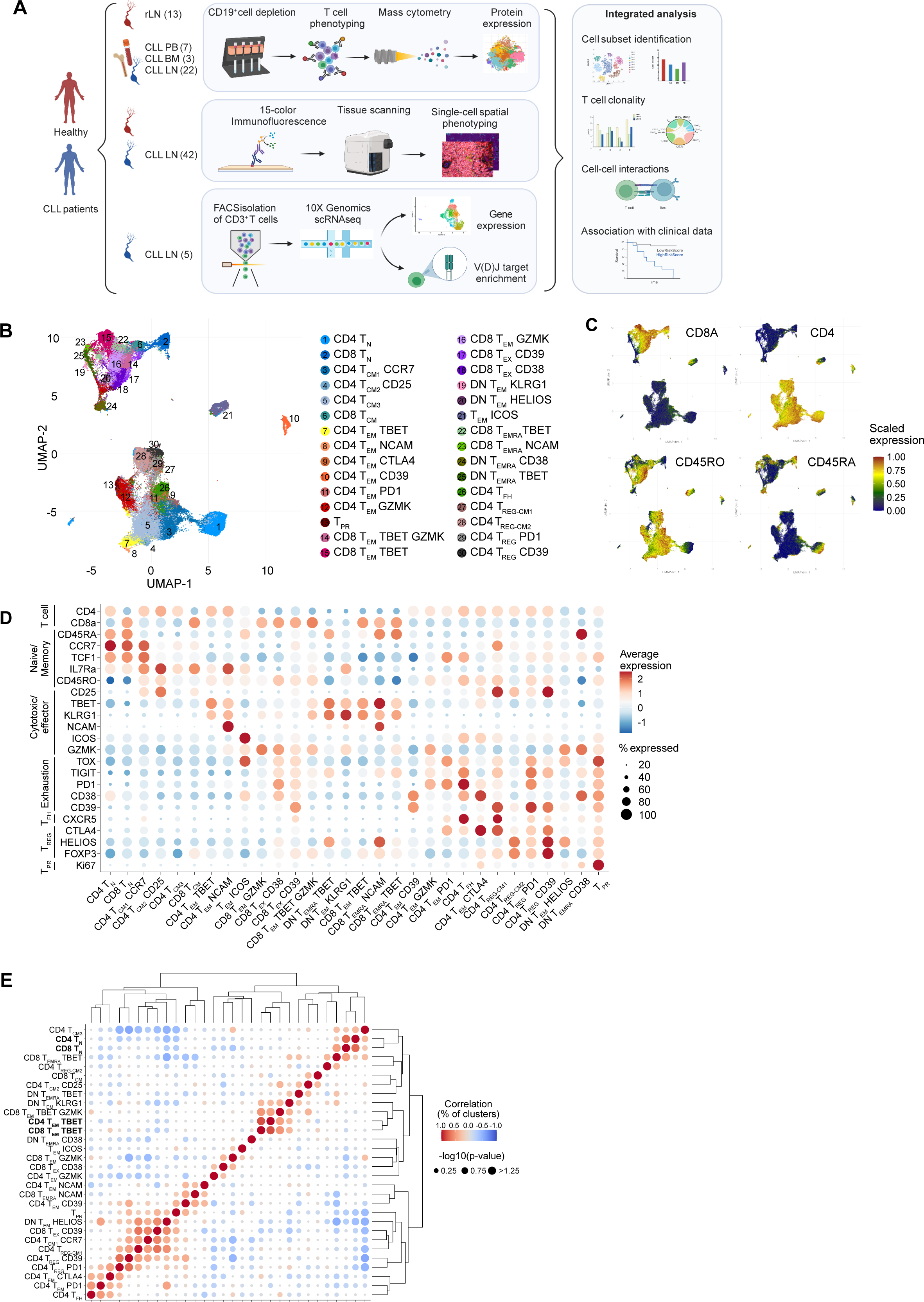
Profiling of the T cell landscape in CLL tissues at the single-cell resolution. **A)** Graphical overview of the study design. Mass cytometry analyses were performed on 13 rLNs, 7 CLL PB, 3 CLL BM and 22 CLL LNs, 15-color multiplex immunostaining was applied in an additional data set of 42 CLL LNs, and 5 CLL LNs were analyzed using scRNA-seq and TCR-seq. **B)** UMAP plot of 5.2 x10^6^ CD3^+^ T cells from 13 rLNs, 7 CLL PB, 3 CLL BM and 22 CLL LNs analyzed by mass cytometry identifying 30 clusters, including 15 for CD4^+^ T cells, 9 for CD8^+^ T cells, 2 clusters containing both CD4^+^ and CD8^+^ T cells, and 4 DN T cells. **C)** Projection of a selection of protein markers identifying T cell states. Cells are colored based on the normalized protein expression. **D)** Dot plot of the expression of marker genes in the 30 cell clusters. **E)** Heatmap showing the Pearson correlation coefficient and its associated p-value of cell subset proportions from the 20 CLL LNs, corresponding to the worst performing subset (for which the p-value was the highest) of all leave-one-out patient sample sets.

To identify relationships between the T cell populations present in the CLL LNs, we correlated the subset frequencies of all individual CLL LN samples (n = 20, excluding the second time point of 2 samples, Figure 1E). Naïve CD4^+^ and CD8^+^ T cells as well as CD4^+^ and CD8^+^ T cells with a SLEC phenotype were significantly associated with each other, indicating that the LN tumor microenvironment (TME) is similarly influencing the differentiation of these cell states both in CD4^+^ and CD8^+^ T cells.

Spatial analyses of immune cell populations by multiplex immunofluorescence staining of a tissue microarray (TMA) of 42 CLL LN samples confirmed the presence and colocalization of malignant B cells with T cells in the tissue (Figure 2A, Suppl. Figure 3A-B). Quantification of the frequencies of the major immune cell subsets in the tissues revealed a positive correlation for CD8^+^ PD1^-^ and CD8 PD1^+^ T cells, for myeloid cells and CD4 T_REG_, and the strongest correlation for CD4 T_REG_ and PD1^+^ CD8^+^ T cells (Figure 2B, Suppl. Figure 3C) which is in line with a role of CD4 T_REG_ in promoting the accumulation of dysfunctional CD8^+^ T cells (Sawant et al., 2019). We next quantitatively assessed the spatial cell-cell interactions by calculating the enrichment of pairwise interacting cell types for all cores using Giotto (Dries et al., 2021). This confirmed that the physical interaction between CD8 PD1^+^ T cells and CD4 T_REG_ as well as CD4 PD1^+^ T cells and CD4 T_REG_ is significantly enriched (Figure 2C-D, Suppl. Figure 3D).

**Figure 2:**
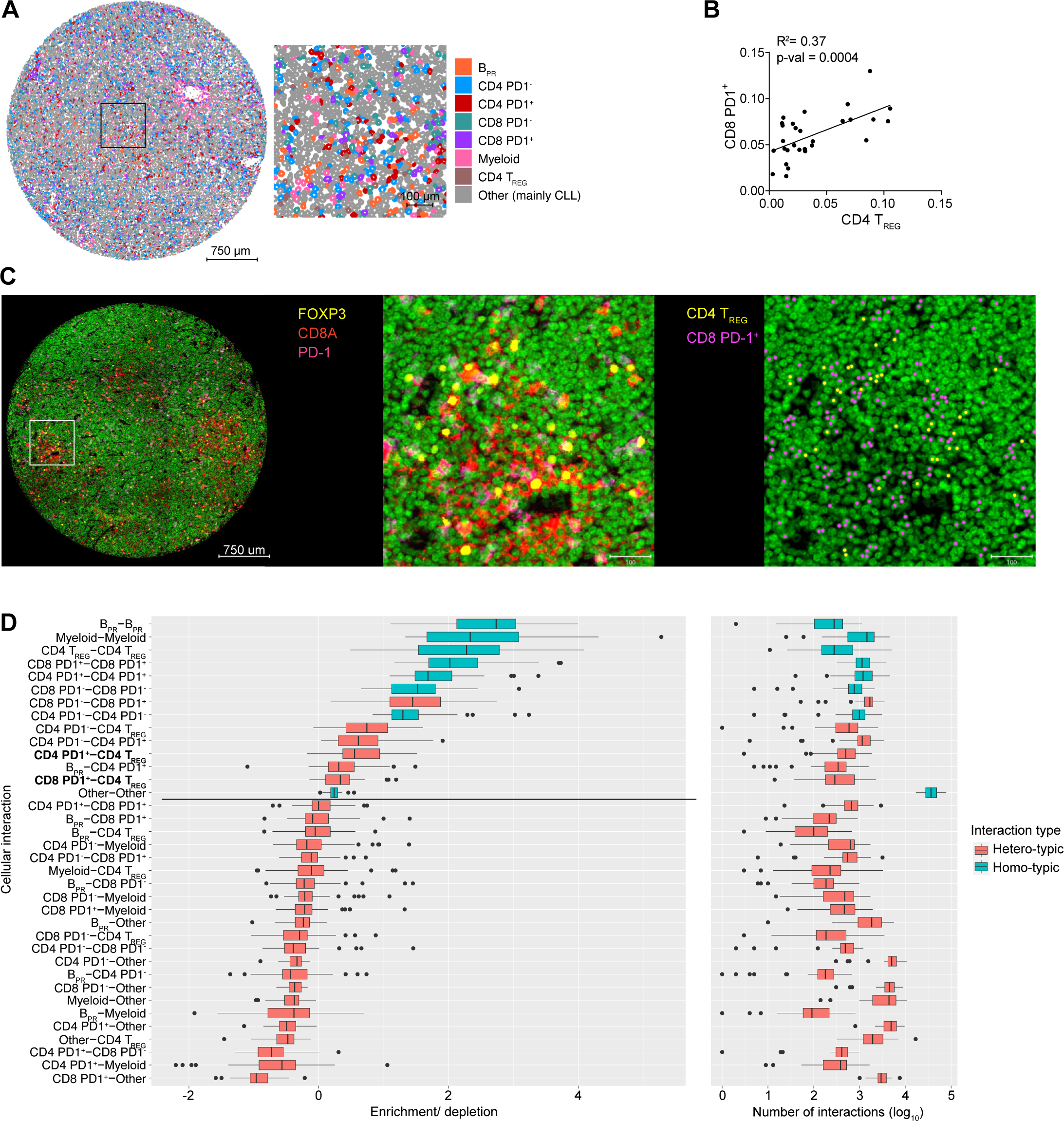
PD-1^+^ T cells reside in close proximity to T_REG_ cells in CLL LNs. **A)** Representative CLL LN-derived tissue core stained with 15-color Orion multiplex with cells colored according to subset identification. CLL cells are displayed as B_PR_: Ki-67^+^ proliferating B cells, and Other: mainly non-proliferating B cells. **B)** Pearson correlations of PD1^+^ CD8^+^ T cells and T_REG_ frequencies determined by multiplex staining of 42 CLL LN cores. Each dot represents an individual patient. **C)** Representative CLL LN-derived tissue core (left image) and field of view (middle image) displaying FOXP3 (yellow), CD8A (red) and PD-1 (pink) staining. The right image displays cells identified as T_REG_ and CD8 PD-1^+^ T cells in yellow and purple dots, respectively. **D)** Boxplots (left) depicting the range of enrichment of pairwise interacting cell types for all cores (n = 42). Boxplots (right) showing the absolute number of interactions in log10 scale between pairwise interacting cell types across all cores. Colored by type of interaction, between same cell type (blue) or different cell types (red).

### CLL LNs represent a distinct niche enriched with immunosuppressive T cell phenotypes

We next examined the distribution of T cell immunophenotypes across the three different tissues analyzed in CLL. The sample composition of PB and BM was similar, but differed from LN samples (Figure 3A-B, Suppl. Figure 4A), suggesting that tissue cues significantly determine T cell composition. PB and BM samples were enriched in SLEC T cells, such as CD8 T_EM_ TBET, CD4 T_EM_ TBET, and DN T_EMRA_ TBET, while CLL LNs contained higher frequencies of exhausted T cells (CD8 T_EX_ CD39 and CD8 T_EX_ CD38), CD4 T_EM_ GZMK cells, CD4 T_FH_ cells, several CD4 T_REG_ subsets (CD4 T_REG_-_CM1_, CD4 T_REG_ PD1, and CD4 T_REG_ CD39), as well as proliferating cells (T_PR_) (Figure 3C). Even though the abundancies among cell clusters within the tissue types correlated, no significant associations of the frequencies of cell subsets between PB and LN samples of the same patient were identified, underscoring a differential composition of T cell subsets between tissues (Suppl. Figure 4B).

**Figure 3:**
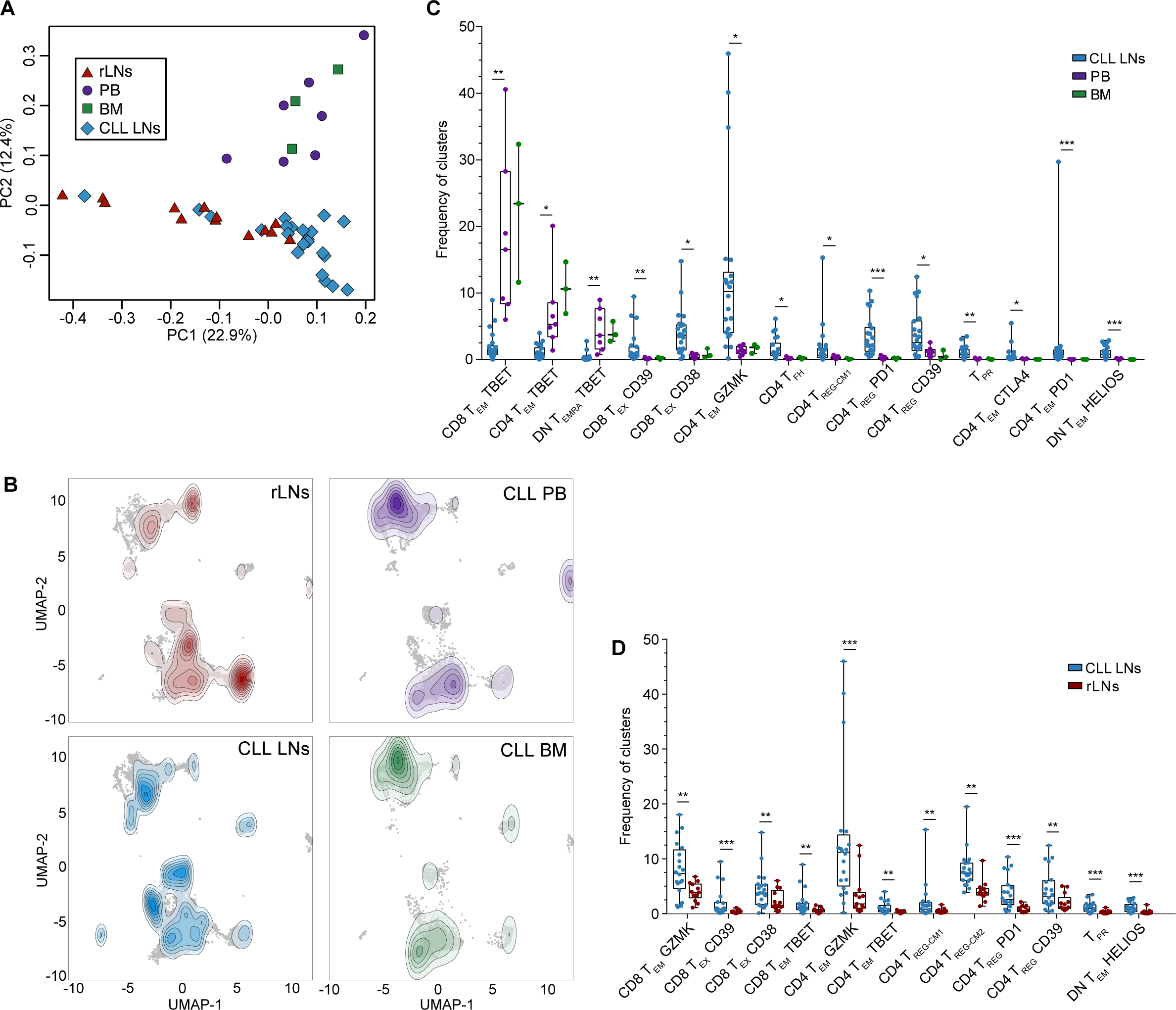
The T cell composition of CLL LNs is distinct and enriched in regulatory and exhausted subsets. **A)** Principal component analysis of all samples analyzed by mass cytometry based on cell subset frequencies. **B)** UMAP plots of T cells from rLNs, CLL PB, CLL BM and CLL LNs overlaid with a contour plot indicating the cell density. **C)** Boxplot showing cell subset abundances in LNs (n = 20), PB (n = 7) and BM (n = 3) of CLL patients. Statistical significance between CLL LNs and CLL PBs was assessed by limma on normalized cell counts. **D)** Boxplot showing cell subset abundances in CLL LNs (n = 20) and rLN (n = 13). Each symbol represents an individual patient sample and statistical significance was tested by limma on normalized cell counts, \**p*<0.05, **<0.01, ***<0.001.

The comparison of cell subset frequencies between CLL LNs and rLNs revealed that higher percentages of cytotoxic CD8 T_EM_ GZMK as well as exhausted CD8^+^ T cells (CD8 T_EX_ CD39 and CD8 T_EX_ CD38) were present in CLL LNs (Figure 3D). Within the CD4 compartment, CD4 T_EM_ GZMK cells and all CD4 T_REG_ clusters (CD4 T_REG-CM1_, CD4 T_REG-CM2_, CD4 T_REG_ PD1, and CD4 T_REG_ CD39) were also enriched in CLL LNs, in addition to proliferating T_PR_ cells and DN T_EM_ HELIOS cells (Figure 3D). Collectively, these results suggest that CLL LNs constitute a distinct niche where malignant B cell accumulation induces an immunosuppressive microenvironment with an enrichment of regulatory T cells and exhausted cytotoxic T cells.

Next, we examined the relation of clinical features to a distinct T cell composition in the CLL LNs. We detected a strong negative association between age of the patients and the frequency of naïve CD8^+^ T cells (Suppl. Figure 4C, Suppl. Table 3) which is in line with published data (Gupta et al., 2004). In addition, the frequencies of 2 CD4 T_REG_ clusters (CD4 T_REG CM2_ and CD4 T_REG_ PD1) showed a positive correlation with age (Suppl. Figure 4D, E). We did not observe significant associations between the frequencies of T cell clusters with sex or the clinical stage of the patients (Suppl. Table 3).

### Definition of T cell exhaustion states in CLL LNs by single-cell RNA-sequencing

Next, single-cell RNA-seq of T cells from CLL LNs was performed to characterize their transcriptional profile and clonal diversity. CD3^+^ T cells and a small spike-in population of CLL cells from LN samples of 5 CLL patients were FACS-sorted and subjected to scRNA-seq using the 10X Genomics platform, yielding 8,825 cells with an average of 4,577 reads per cell and a median of 1,475 genes per cell (STAR methods, Suppl. Figure 5A). Using unsupervised graph-based clustering, we identified 13 clusters presented in a UMAP plot (Figure 4A, STAR methods), for which we assigned an identity based on the differential gene expression of each cluster compared to the rest of the cells (Figure 4B). All cell clusters were shared among patients although at different proportions (Figure 4C, Suppl. Figure 5B). Clusters were defined as CLL cells, a small cluster of dendritic cells (DCs), 6 CD4^+^ T cell subsets, 4 CD8^+^ T cell subsets, and a KI67^+^ proliferating T cell cluster (T_PR_) which contained both CD4^+^ and CD8^+^ T cells (Figure 4A, B). More specifically, both CD4^+^ and CD8^+^ naïve (CD4 T_N_ and CD8 T_N_) T cells were defined based on the expression of marker genes such as *SELL*, *CCR7* and *IL7R*; regulatory CD4^+^ T cells (CD4 T_REG_) were identified by high expression of *FOXP3*, *IL2RA* and *IKZF2*; a heterogeneous cluster of cells containing mainly follicular helper CD4^+^ T cells (CD4 T_FH_) was defined by the expression of *CXCR5*, *CD200*, *TOX2* and *TOX* (Suppl. Figure 5C, D), the small proportion of CD8^+^ T cells in this cluster is in line with a shared transcriptional program of exhausted CD8^+^ T cells with T_FH_ (Im et al., 2016). Conventional CD4^+^ T cells with a central memory phenotype were identified by the high expression of *IL7R, CD40LG, KLRB1* as well as *CD69* (CD4 T_H_ CD69) or *ITGB1* and *KLF2* (CD4 T_H_ KLF2) in accordance with published data of other B cell lymphoma (Roider et al., 2022). A heterogeneous cluster mainly comprising CD4^+^ T cells expressed memory marker and effector molecule genes such as *GZMK*, resembling the CD4 T_EM_ GZMK subset that shares features with T_R_1-like cells, identified by mass cytometry (Figure 1B). CD8^+^ effector memory subsets were characterized by a higher expression of effector molecule genes like *CCL5, NKG7, PRF1*, *GZMA* and *GZMK*. One subset expressed generally lower levels of these genes and high levels of *LYAR* (CD8 T_EM_), another expressed higher levels of *CCL5* (CD8 T_PEX_), and resembled the CD8 T_EM_ GZMK cluster identified by mass cytometry (Figure 1B), and a third showed elevated expression of genes related to exhaustion such as *HAVCR2* (coding for TIM3), *LAG3, ENTPD1* (CD39), *PDCD1* (PD1), *CTLA4* and *TIGIT* (CD8 T_EX_), and therefore resembled the two T_EX_ clusters defined in the CyTOF data by high expression of these exhaustion markers (Figure 1B). Notably, the later subset expressed highest levels of the exhaustion signature from Zheng *et al*. (Zheng et al., 2021) (Figure 4D) confirming its dysfunctional state, while CD8 T_PEX_ cells presented the highest score of the precursor exhausted signatures from Guo *et al*. (Guo et al., 2018) (Figure 4E) and Andreatta M *et al*. (Andreatta et al., 2021) (Suppl. Figure 5E) suggesting that these cells constitute a precursor state of exhausted CD8^+^ T cells commonly named T_PEX_ cells (Kallies et al., 2020). Subset clustering and definition were in agreement with the annotation from the pbmc3k signatures from Hao *et al*. (Hao et al., 2021) supporting the validity of our annotations (Suppl. Figure 5F). In addition, the frequency of the main T cell clusters was comparable and positively correlated between scRNA-seq and mass cytometry in the 4 samples analyzed by both techniques (Suppl. Figure 5G, H), underscoring the robustness of our data.

**Figure 4:**
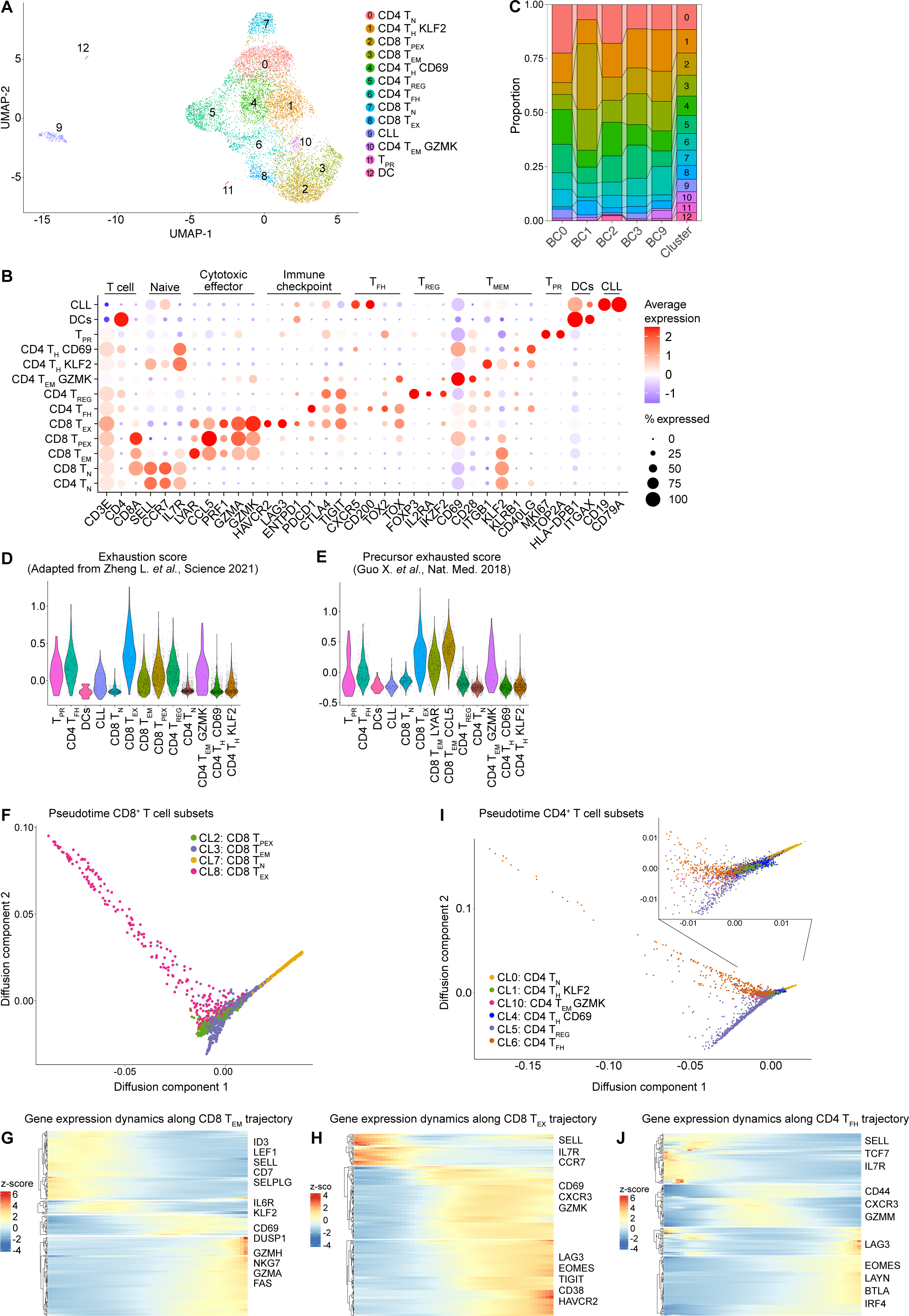
Single-cell RNA-seq defines T cell states in CLL LNs. **A)** UMAP plot of 8,576 cells from 5 CLL LNs analyzed by scRNA-seq identifying 13 clusters, including 6 for CD4^+^ T cells, 4 CD8^+^ T cells, proliferating CD4^+^ and CD8^+^ T cells, DCs and spiked in CLL cells. **B)** Dot plot of the expression of marker genes in the 13 cell clusters. **C)** Cell subset abundance in each sample (BC0, 1, 2, 3, and 9), with colors and numbers corresponding to the identified cell clusters. **D-E)** Violin plot of average expression levels of the exhaustion gene signature adapted from Zheng L, *et al* Science, 2021 (D), and the precursor exhaustion gene signature from Guo X, *et al* Nat Med, 2018 (E). **F)** Pseudotime trajectory across the 4 CD8^+^ T cell subsets**. G)** Heatmap showing genes with significant expression changes along the trajectory from CD8 T_N_ to CD8 T_EM_ **H)** Heatmap showing genes with significant expression changes along the trajectory from T_N_ to CD8 T_EX_. Color represents z-scores. **I)** Pseudotime trajectory across the 6 CD4^+^ T cell subsets**. J)** Heatmap showing genes with significant expression changes along the trajectory from CD4 T_N_ to CD4 T_FH_. Color represents z-scores.

To delineate the relationship between T cell states, we performed trajectory analyses of CD4^+^ and CD8^+^ T cells by destiny (Angerer et al., 2016). When defining CD8 T_N_ cells as the starting point of the differentiation path, we observed a bifurcation into two branches, one containing CD8 T_EM_ cells and the other CD8 T_PEX_ cells, from which T_EX_ cells diverged and constituted a terminal state (Figure 4F, Suppl. Figure 5I). When inspecting the differentiation path from CD8 T_N_ to CD8 T_EM_, four clusters of differentially expressed genes (DEGs) were identified along the trajectory. Naïve marker genes (*LEF1, SELL*) were highly expressed in the first cluster, the second included the transcription factor *KLF2* and *IL6R*, the third activation marker genes (*CD69* and *DUSP1*), and the fourth effector molecule genes (*GZMH, NKG7, GZMA and FAS*) (Figure 4G). The differentiation path from CD8 T_N_ to CD8 T_PEX_ and CD8 T_EX_ was also comprised of four clusters of DEGs, with naïve marker genes (*SELL*, *IL7R* and *CCR7*) expressed at the beginning, followed by genes associated with activation (such as *CD69*) and cytotoxic function (*GZMK*), and genes related to exhaustion (*LAG3, EOMES, TIGIT, CD38* and *HAVCR2*) that were highest at the end of the trajectory (Figure 4H). This data suggests GZMK^+^ CD8^+^ T cells as a precursor state of terminally exhausted CD8^+^ T cells which is in line with previously published data (Liu et al., 2022). Together with our CyTOF results that showed an accumulation of the CD8 T_EM_ GZMK precursor state and terminally exhausted CD8^+^ T cells in CLL LNs (Figure 3D), these findings suggest cancer-driven exhaustion of CD8^+^ T cells in the lymphatic tissue of patients with CLL.

The CD4^+^ T cell trajectory followed a similar pattern as CD8^+^ T cells, starting with naïve genes (*SELL, TCF7* and *IL7R*), followed by genes for activation and effector molecules (*CD44, CXCR3,* and *GZMM*), and then by exhaustion-related genes (*LAG3, EOMES, LAYN, BTLA,* and *IRF4*) (Figure 4I, J; Suppl. Figure 5J), indicating a cancer-driven exhaustion also of CD4^+^ T cells. The CD4 T_FH_ and CD4 T_REG_ cell clusters were diverting into two branches of the trajectory already at an early stage (Figure 4I).

### Effector memory CD8 T cells resembling precursors of exhausted cells are clonally expanded in CLL LNs

The reconstruction of the T cell receptor (TCR) sequences at the single-cell level allowed us to examine the clonal diversity of T cells within the identified subsets. T cells that shared the same TCR sequence with at least one other cell were defined as clonal, and clone sizes of small (2-5 cells), medium (6-20 cells), large (21-100 cells) and hyperexpanded (101-500 cells) were identified. This data showed a major clonal expansion of CD8^+^ T cells (ranging from 18% to 69%), but not CD4^+^ T cells (4% to 26%) in the 5 CLL LN samples, and no correlation between the expansion rate of CD4^+^ and CD8^+^ T cells (Figure 5A). A comparison of the TCR sequences of the major clones with VDJdb verified that none of these TCRs were linked to known viral antigens (Suppl. Table 4). The clonal composition within the CD8^+^ T cell compartment was highly variable among patients, with most samples showing mostly small and medium sized clones, and one sample (BC1) presenting one strongly dominating, hyperexpanded clone. Notably, CD8^+^ T cell clonal expansion was enriched in the two CD8 T_EM_ subsets and in CD8 T_EX_ cells (Figure 5B, C) which explains the relative enrichment of the CD8 T_EM_ cluster in the BC1 sample (Figure 4C) and suggests that these cells are tumor-reactive. Most TCR clones were confined to a specific T cell cluster, but some were shared between clusters with similar functional phenotypes, such as CD8 T_EM_ and CD8 T_PEX_ (Figure 5D, Suppl. Figure 6A). This suggests that either a dividing activated T cell produces cells with distinct transcriptional states, or that cells dynamically transit between two transcriptional states.

**Figure 5:**
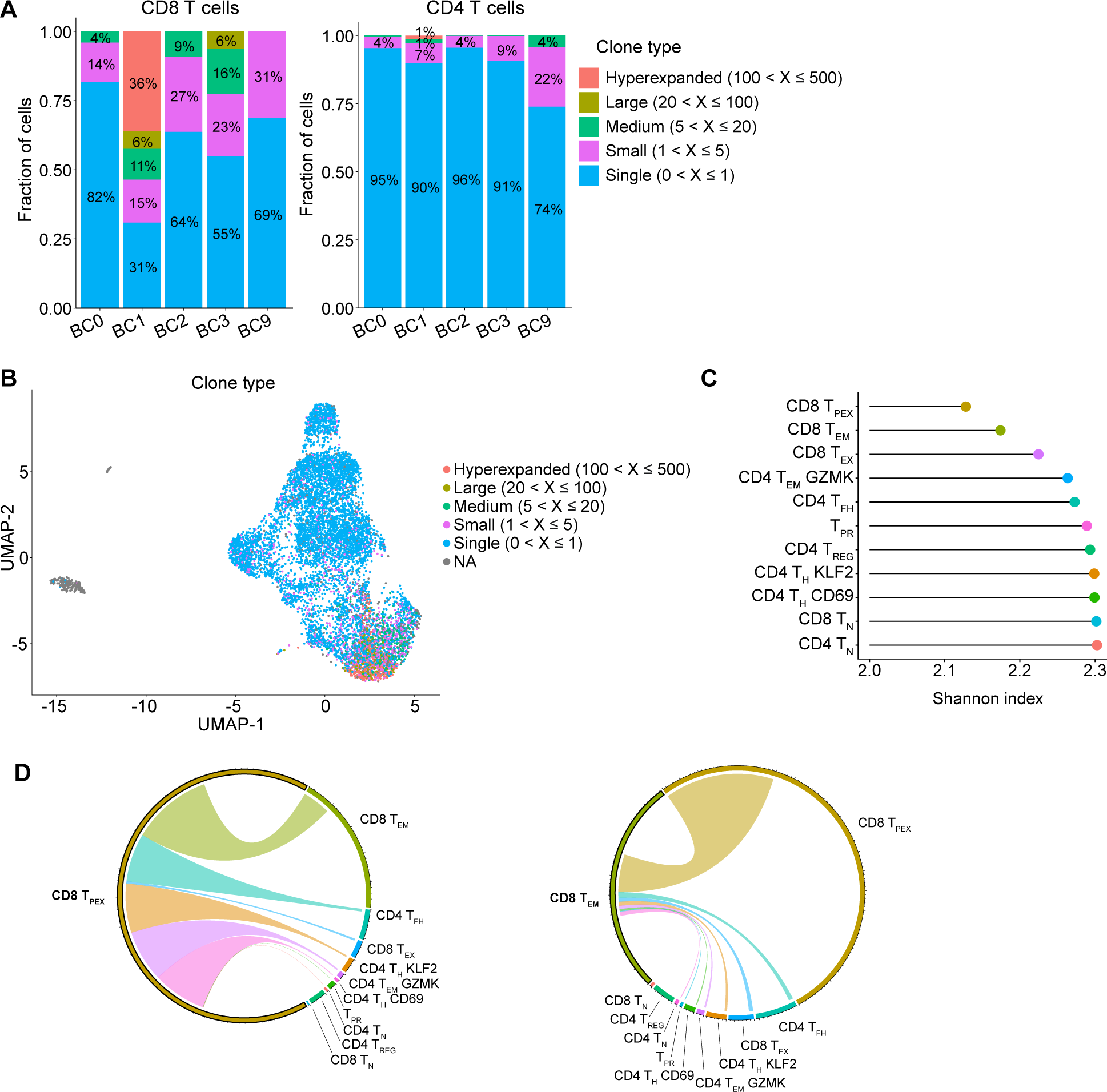
T cell clonal expansion in CLL is mainly restricted to CD8 T_EM_ cells. **A)** Bar plot indicating the percentage (rounded values are indicated) of single, small, medium, large and hyperexpanded sized clones in CD8^+^ (left) and CD4^+^ (right) T cells for each patient analyzed (n = 5). **B)** UMAP plot colored according to the T cell clone size based on the TCR-seq data. **C)** Graph showing the TCR Shannon diversity index for each T cell subset identified by scRNA-seq. The dot color corresponds to the UMAP cluster plot from Figure 4A. **D)** Chord diagram showing clone sharing from CD8 T_EM_ and CD8 T_PEX_. Only cells carrying expanded TCR sequence are presented in the circle. Cells sharing identical TCR sequences across clusters are summed and connected by bands.

### CLL cells and T cells establish a unique interactome including Galectin-9 and TIM3 as ligand-receptor pair

Next, we predicted receptor-ligand interactions between T cells and CLL cells in the LNs by performing differential interaction analysis using CellChat (Jin et al., 2021) on the basis of the public repository of ligand-receptor interactions from OmniPath (Türei et al., 2016). We included published scRNA-seq data from 5 rLNs (Aoki et al., 2020) in this analysis (Suppl. Figure 7A, B) to predict CLL-specific interactions. T cell subsets in both datasets were grouped into major functional types (CD4 T_N_, CD8 T_N_, CD4 T_FH_, CD4 T_CM_, CD4 T_REG_, CD4 T_EM_ and CD8 T_EM_) to facilitate comparison. Predicted receptor-ligand interactions were prioritized by the probability of differential interaction across CLL LNs and rLNs. Overall, the number of inferred interactions was slightly higher in the CLL LNs compared to the rLNs (1828 vs. 1622; Suppl. Figure 7C). The highest number of predicted interactions was found for CLL cells and CD8 T_EM_ cells, which act as senders and receivers of signals, both in an autocrine and paracrine manner (Figure 6A, B). These two cell types were predicted to strongly interact with each other and also with all other cell subsets. Inferred CLL-specific interactions included ligand-receptor pairs with known relevance in CLL, such as CD74-MIF (Reinart et al., 2013), and the chemokines CCL4 and CCL5 and their receptors (Burger et al., 2009; Kiaii et al., 2013) (Figure 6C). In addition, collagen and adhesion molecules and their binding partners (COL9A2/3 and ICAM-ITGAL) were predicted as novel interactions. Numerous inferred interactions between T cells and CLL cells were related to T cell inhibitory signals, such as HLA-LAG3 (Wierz et al., 2018), BTLA-CD247 (Karabon et al., 2020) and LGALS9-related circuits. Galectin-9 (encoded by *LGALS9*) is a known ligand of the inhibitory receptor TIM3 and its binding to TIM3 induces T cell death and thus contributes to tumor immune escape (Yang et al., 2008; Zhu et al., 2005). In CLL, Galectin-9 levels are increased in serum of patients (Pang et al., 2021b; Wdowiak et al., 2019; Xierenguli et al., 2020). Our scRNA-seq data identified CLL cells and dendritic cells as a source of Galectin-9, showing an increased *LGALS9* expression in CLL compared to rLNs (Figure 6D). The Galectin-9 binding partner TIM3 (encoded by *HAVCR2*) was detected in CD8 T_EX_ cells in CLL LNs, but was absent in rLNs (Figure 6D). Multiplex immunofluorescence staining of CLL LN sections further confirmed the presence of TIM3 expressing CD4^+^ and CD8^+^ T cells in the tissue (Figure 6E).

**Figure 6:**
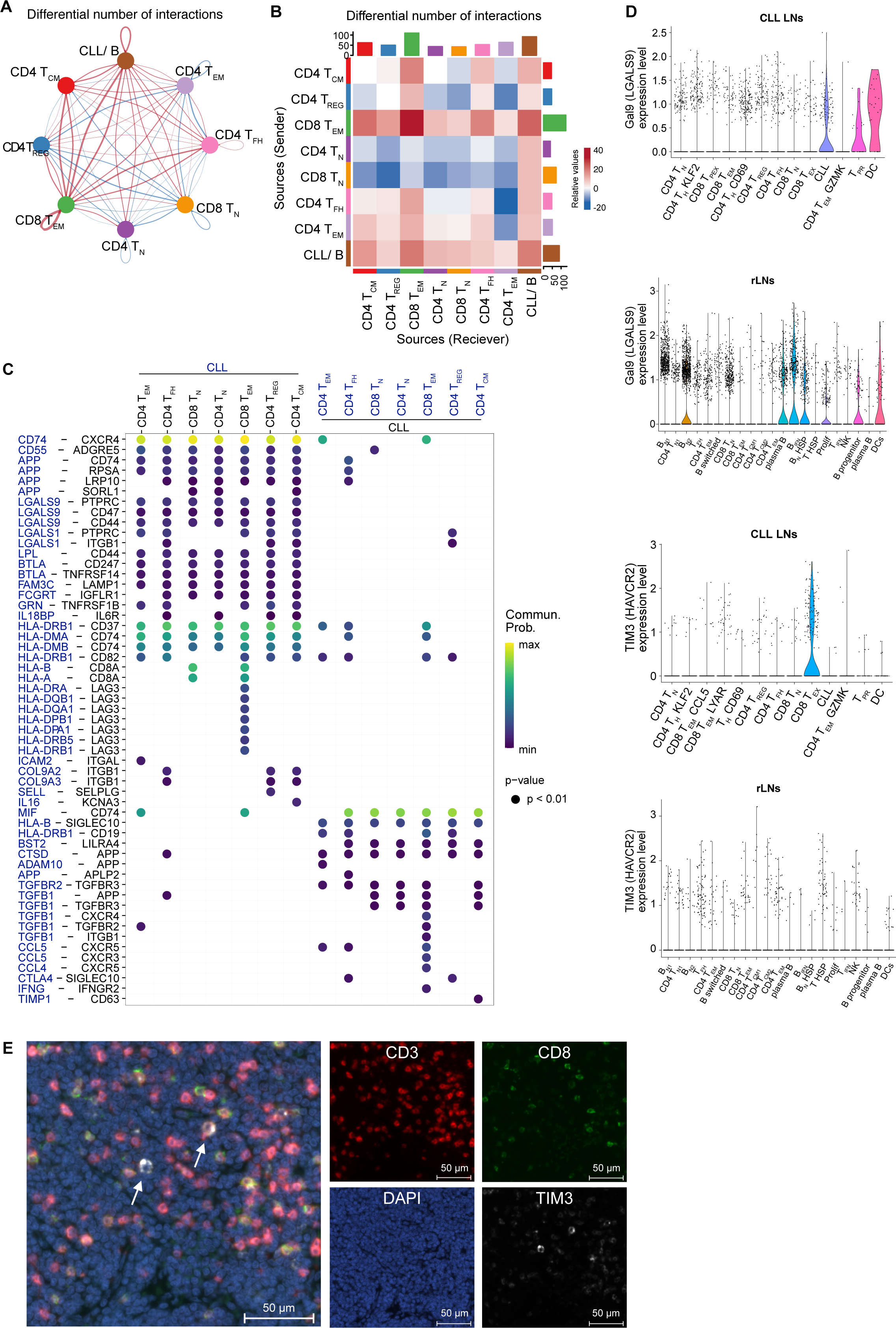
Interactome analyses of CLL LNs identify Galectin-9 as a potential therapeutic target. **A)** Circle plot depicting the differential number of interactions between cell subsets in CLL LNs compared to rLNs. Thickness of bands represents the number of differential interactions between the two data sets, and increased interactions are depicted in red, decreased interactions in blue. **B)** Heatmap showing the differential number of interactions of CLL LNs versus rLNs. Rows and columns represent source and target subsets, respectively. Bar plots represent the total outgoing (right) and incoming (top) interactions scores respectively. **C)** Heatmap plot depicting a curated list of ligand-receptor pairs differentially upregulated in CLL LNs compared to rLNs as identified via CellChat (Jin et al., 2021). The dot color and size represent the calculated communication probability and p-values of differential communication, respectively. The first 7 columns show interactions sent by CLL cells, while the last 7 columns show interactions received by CLL cells. **D)** Violin plots showing the expression distribution of *LGALS9* (top two plots) and *HAVCR2* (bottom plots) genes in the CLL LN data set and the rLN data set from Aoki *et* al. (Aoki et al., 2020). Each dot represents one cell. **E)** Representative microscopy images showing CD3, CD8, TIM3 and DAPI as single stains and overlayed image (left) in a CLL LN tissue section. Arrows indicate CD3^+^ CD8^+^ TIM3^+^ cells.

### Galectin-9 blockade reduces CLL development in mice

We next aimed to preclinically test the potential of Galectin-9 blockade in a mouse model of CLL. As the Eµ-TCL1 mouse line (Bichi et al., 2002) is the most explored and used *in vivo* model of CLL, we first performed single-cell RNA-seq to characterize the T cell compartment of mice that developed CLL-like disease after adoptive transfer of malignant B cells from this mouse line (TCL1 AT). This allowed us to identify 13 clusters of T cells in the spleen of leukemic mice, including naïve, effector, effector memory, and regulatory T cell subsets (Suppl. Figure 8A, B, C). When integrating TCR-seq data, it became clear that one of the two analyzed samples (#107) harbored a hyperexpanded CD4^+^ T cell clone showing an effector phenotype and *Gzmk* expression, which was not the case for the second sample (#110) that contained a large CD8^+^ effector memory T cell cluster (Suppl. Figure 8D, E). The clonally expanded CD4^+^ effector T cells in sample #107 contained a cluster of cells expressing exhaustion-related genes. This data suggests the occurrence of CD4^+^ T cell-driven adaptive immune control in TCL1 AT mice which is in line with observations in other tumor mouse models and patients with cancer (Oh and Fong, 2021; Oh et al., 2020; Quezada et al., 2010; Xie et al., 2010).

Flow cytometry analyses showed that the vast proportion of malignant B cells in the spleen and bone marrow of TCL1 AT mice expressed Galectin-9, which was not the case for B cells from wild-type (WT) mice (Figure 7A, Suppl. Figure 8F). Also, higher frequencies of Galectin-9-positive CLL cells were detected in lymph nodes and PB of TCL1 AT mice compared to WT mice (Suppl. Figure 8F). We further observed TIM3-positive T cells in tumor-bearing TCL1 AT but not WT mice, with the highest frequencies detected in CD8 T_EF_ and CD4 T_REG_ cells (Figure 7B).

**Figure 7:**
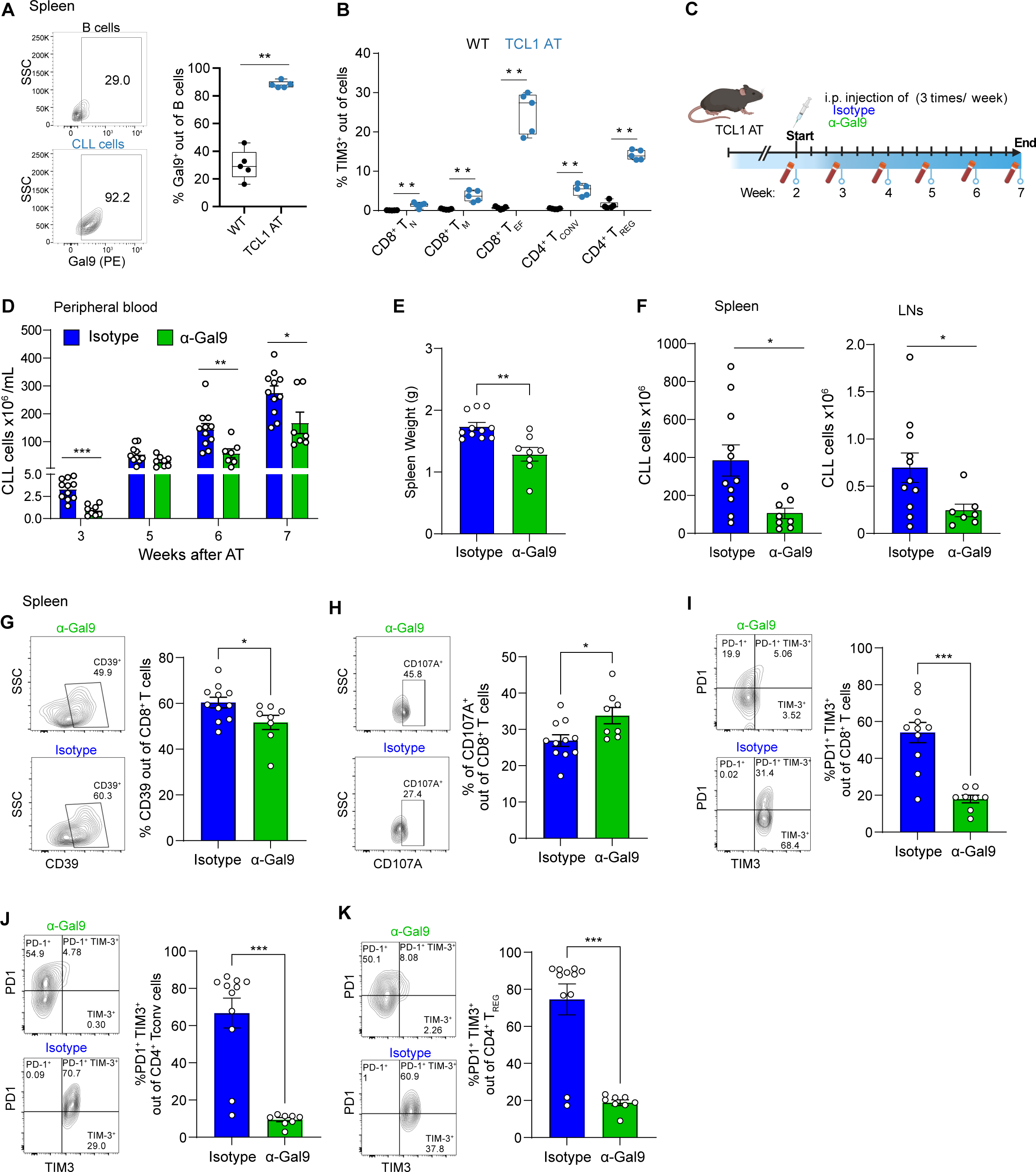
Blocking of Galectin-9 controls tumor growth in the TCL1 mouse model. **A)** Representative contour plot and percentage of Galectin-9^+^ B cells and CLL cells from spleen of WT and TCL1 AT mice, respectively (n = 5). **B)** Percentage of TIM3^+^ cells out of CD8 T_N_, CD8 T_M_, CD8 T_EF_, CD4 T_CONV_ and T_REG_ T cells in spleen of WT and TCL1 AT mice (n = 5). **C)** Schematic diagram of Galectin-9-blocking antibody treatment in TCL1 AT mice (n = 8-10). Analyses of T, myeloid and CLL cells were performed 7 weeks after treatment start. **D)** Absolute number of CD19^+^ CD5^+^ CLL cells in blood of isotype antibody- and anti-Galectin-9-treated mice. **E)** Spleen weight, and **F)** absolute number of CD19^+^ CD5^+^ CLL cells in spleen and lymph nodes of isotype antibody- and anti-Galectin-9-treated mice. **G)** Representative contour plot and percentage of CD39^+^ cells out of CD8^+^ T cells from spleen of isotype antibody- and anti-Galectin-9-treated mice. **H)** Representative contour plot and percentage of CD107A^+^ out of CD8^+^ T cells from spleen of isotype antibody- and anti-Galectin-9-treated mice. **I-K)** Representative contour plot and percentage of PD1^+^ TIM3^+^ cells out of CD8^+^ T cells (I), CD4^+^ T_CONV_ (J), and T_REG_ (K) cells from spleen of isotype antibody- and anti-Galectin-9-treated mice. Each symbol represents an individual mouse, statistical significance was tested by unpaired t-test with Welch approximation, \**p*<0.05, **<0.01, ***<0.001. Bars indicate mean ± SEM.

To test the potential of Galectin-9 as novel therapy target for CLL, we treated CLL-bearing TCL1 AT mice with Galectin-9-blocking antibodies for 20 days (Figure 7C). By a weekly assessment of CLL cell counts in the blood of the mice, a deceleration of leukemia development was observed by anti-Galectin-9 compared to isotype control treatment (Figure 7D). At the endpoint of the experiment, 7 weeks after TCL1 AT, a reduced spleen weight and lower numbers of CLL cells were detected in the spleen and lymph nodes of the mice receiving anti-Galectin-9 (Figure 7E, F). We further analyzed T cells in the spleen and lymph nodes at the endpoint and did not observe major differences in the numbers of CD8^+^ T cells, conventional CD4^+^ T cells, or CD4^+^ T_REG_, nor in their activation state measured as % of CD44^+^ cells (Suppl. Figure 9A, B, C). We detected however, a lower frequency of CD39^+^ and a higher frequency of CD107A^+^ CD8^+^ T cells in the treated mice (Figure 7G, H), suggesting a higher functionality of CD8^+^ T cells upon anti-Galectin-9 treatment. The most drastic differences were observed in the frequencies of TIM3^+^ T cells, which were severely decreased for CD8^+^ T cells, conventional CD4^+^ T cells and CD4^+^ PD1^+^ T_REG_ cells in the spleen and to a lower degree also in LNs (Figure 7I-K, Suppl. Figure 9D) which is in line with the proposed role of Galectin-9 in preventing apoptosis of TIM3^+^ T cells (Zhu et al., 2005). Based on these data, we suggest that anti-Galectin-9 treatment results in a better immune control of CLL via suspending the immune suppressive activity of TIM3 on CLL-associated T cells.

### High Galectin-9 expression is associated with shorter survival of patients with CLL, kidney cancer and brain tumors

Next, we explored the expression of *LGALS9* across cancer entities using publically available TCGA data (Li et al., 2021). A comparison of tumor and respective normal tissue revealed a higher expression in about half of the cancer types analyzed (Figure 8A, Suppl. Figure 10A). To estimate the relevance of Galectin-9 in CLL development, we analyzed a previously published proteome data set of CLL cells (Herbst et al., 2022). Dividing patients into two groups based on the median level of Galectin-9 expression clearly demonstrated a significantly shorter treatment-free survival in cases with high Galectin-9 expression (Figure 8B). Dividing this cohort of patients into the two main prognostic groups, namely cases with unmutated or mutated *IGHV* gene locus, we observed higher Galectin-9 protein expression in the *IGHV*-unmutated group which has a worse overall prognosis (Suppl. Figure 10B). Within this group, higher Galectin-9 protein levels clearly predicted shorter treatment-free survival (Suppl. Figure 10C) which was not the case in the *IGHV*-mutated group of CLL patients (Suppl. Figure 10D).

**Figure 8:**
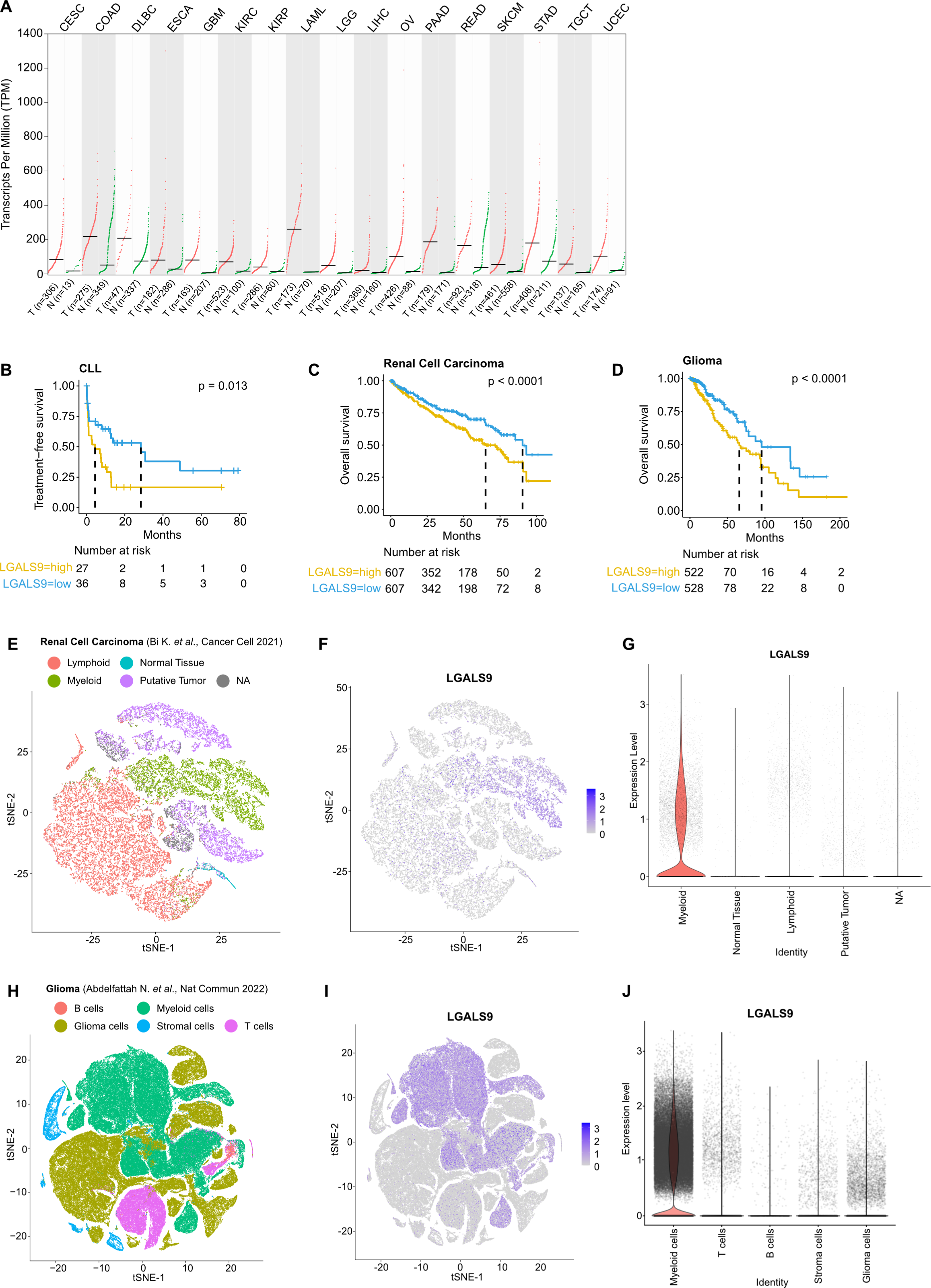
Elevated *LGALS9* expression correlates with poor survival in cancer patients. **A)** *LGALS9* transcript levels in tumor (T) versus healthy (N) tissue in CESC (Cervical squamous cell carcinoma and endocervical adenocarcinoma), COAD (Colon adenocarcinoma), DLCL (Diffuse large B cell lymphoma), ESCA (Esophageal carcinoma), GBM (Glioblastoma multiforme), KIRC (Kidney renal clear cell carcinoma), KIRP (Kidney renal papillary cell carcinoma), LAML (Acute Myeloid Leukemia), LGG (Brain Lower Grade Glioma), LIHC (Liver hepatocellular carcinoma), OV (Ovarian serous cystadenocarcinoma), PAAD (Pancreatic adenocarcinoma), READ (Rectum adenocarcinoma), SKCM (Skin Cutaneous Melanoma), STAD (Stomach adenocarcinoma), TGCT (Testicular Germ Cell Tumors), and UCEC (Uterine Corpus Endometrial Carcinoma) (Li et al., 2021). **B)** Time-to-treatment in CLL patients with high or low Galectin-9 protein levels (n = 63) (Herbst et al., 2022). **C-D)** Overall survival in renal cell carcinoma (C) and glioma (D) patients with high or low *LGALS9* transcript levels. **E-J)** Single-cell RNA-seq analysis of tumor samples in renal cell carcinoma (Bi et al., 2021) (E-G) and glioma (Abdelfattah et al., 2022) (H-J). E and H) UMAP plots identifying tumor, infiltrating immune cells and normal tissue cells from E) 8 renal cell carcinoma patients, and H) 9 glioma patients. F and I) UMAP plot displaying *LGALS9* expression. G and J) Violin plots of *LGALS9* expression.

We further analyzed the prognostic value of *LGALS9* expression across cancer types and identified significant associations of high levels of *LGALS9* with shorter overall survival for renal cell carcinoma and glioma patients (Figure 8C+D). By exploring published scRNA-seq data of these two cancer types, we observed *LGALS9* expression mainly in tumor-associated myeloid cells (Figure 8E-J). Of interest, myeloid cell infiltration has been linked to immune suppression and worse outcome of patients for renal cell carcinoma and glioma (Jikuya et al., 2020; Şenbabaoğlu et al., 2016; Vuong et al., 2019; Zhang et al., 2021). Altogether this suggests Galectin-9 as novel potential target for immunotherapy in CLL and likely other cancer entities.

## Discussion

T cell exhaustion is considered as one of the main reasons for failure of immunotherapies. However, its contribution to resistance of patients with B cell malignancies to such treatments has not been explored so far. Our study provides, for the first time, a single-cell-resolved analysis of the T cell landscape in blood and tissue samples of CLL, the most common leukemia in the Western world. CLL is a disease that is dependent on microenvironmental stimuli provided by the lymph node niche that drive growth and survival of malignant cells which are quiescent in peripheral blood. Most previous studies characterized disease-associated alterations in the T cell compartment in blood samples. Our data now clearly show that exhaustion of cytotoxic T cells and accumulation of regulatory T cells is very prominent in the lymph nodes but not in blood, suggesting the lymphoid tissue as the site of cancer-directed immune responses in CLL.

Our multimodal omics approach allowed us to integrate scRNA-seq, CyTOF, and multiplex imaging of lymph nodes to obtain an unprecedented resolution of the T cell landscape in CLL. The three complementary data sets enabled the identification and characterization of different stages of T cell exhaustion and their clonal expansion in CLL lymph nodes, as well as several regulatory T cell clusters that are in terms of abundance and spatial localization linked to PD1^+^ CD8^+^ effector T cells. In addition to terminally exhausted CD8 T cells, called T_EX_, the lymph nodes contain a population of cells that resembled the previously described precursor cells of exhaustion, (Im et al., 2016; Utzschneider et al., 2016), now frequently named T_PEX_ (Kallies et al., 2020), that share features with memory T cells and have a high *GZMK* expression (Liu et al., 2022). These cells have self-renewal capacity which likely explains our observation of a strong accumulation of TCR clonotypes in this population. Our data identifying such cells in CLL LNs is in line with recent studies demonstrating their presence in tumor-draining LNs of solid cancer where they are essential for efficient response to ICI therapy (Connolly et al., 2021; Huang et al., 2022).

CLL cells in lymph nodes are densely packed together with T cells and interactome analyses using the scRNA-seq data suggested a list of known and novel molecular interactions between the two cell types. It is well-accepted that CLL cells depend on microenvironment-derived signals for survival and proliferation, and the identified interactions including MIF-CD74 (Reinart et al., 2013), CCL4 and CCL5 (Kiaii et al., 2013), and INFG (Buschle et al., 1993) are part of this support. Further, the interactome study inferred several inhibitory molecular interactions including BTLA-CD247, CTLA4 and Galectin-9 (encoded by *LGALS9*). Galectin-9 is a known ligand of TIM3, an inhibitory receptor that is expressed on exhausted T cells (Wolf et al., 2020). Even though we detected very low levels of *HAVCR2* (encoding TIM3) by scRNA-seq, we confirmed the presence of TIM3-positive T cells in CLL lymph node tissue, and assumed an involvement of this interaction in immune escape as Galectin-9 binding to TIM3 promotes CD4^+^ T_REG_ development in CLL (Pang et al., 2021a).

TIM3-targeting antibodies now enter clinical trials and show the first promising anti-tumor activity in patients with advanced solid tumors (Curigliano et al., 2021). These antibodies block the interaction of TIM3 with phosphatidylserine and CEACAM1, but only partially the binding to Galectin-9 (Sabatos-Peyton et al., 2020; Sabatos-Peyton et al., 2018). Blocking antibodies targeting Galectin-9 can overcome this limitation and suppress tumor growth in combination with chemotherapy in a breast cancer mouse model (de Mingo Pulido et al., 2018). This activity was dependent on CD8^+^ T cells, which is in line with our observation of an enhanced CD8^+^ effector function in anti-Galectin-9 treated TCL1 mice.

Binding of Galectin-9 to TIM3 on T cells induces cell death, thereby limiting adaptive immunity (Zhu et al., 2005). This raises the question of why terminally exhausted T cells harboring TIM3 expression persist in the tumor microenvironment. Recent data showed that PD1, via physically interacting with Galectin-9 and TIM3, protects T cells from apoptosis (Yang et al., 2021). As a consequence, PD1^+^ TIM3^+^ T cells accumulate in tumors even in the presence of enhanced levels of Galectin-9. Blockade of Galectin-9 *in vitro* and in mouse models results in enhanced T cell survival and improved anti-tumor immunity, especially in combination with co-stimulatory anti-GITR treatment (Yang et al., 2021). In accordance, anti-Galectin-9 treatment of TCL1 mice diminished PD1^+^ TIM3^+^ T cells along with a reduction in tumor development.

Besides improving T cell responses, antibody-mediated targeting of Galectin-9 on B-ALL cells induces DNA damage, alters cell cycle progression, and promotes apoptosis *in vitro* and significantly extends the survival of mice with aggressive B-ALL (Lee et al., 2022). Accordingly, we cannot exclude that a similar direct activity on CLL cells contributes to the delayed leukemia development in TCL1 mice. Further, ligand activation of Galectin-9 in the tumor microenvironment of pancreatic ductal adenocarcinoma results in tolerogenic macrophage programming and adaptive immune suppression (Daley et al., 2017). As CLL is dominated by immunosuppressive myeloid cells (Hanna et al., 2019), Galectin-9 blockade potentially ameliorates this milieu which likely contributes to a better T cell function.

In line with these mechanistic findings, *LGALS9* expression is higher in tumor compared to respective normal tissue across many cancer types. In CLL, we defined the malignant B cells as the main source of Galectin-9, even though myeloid cell types likely contribute to its expression. In solid cancers however, Galection-9 is likely produced mainly by myeloid cells, like tumor-associated macrophages or myeloid-derived suppressor cells. This also explains that both the abundance of tumor-infiltrating myeloid cells and high expression of *LGALS9* predict shorter survival of patients with renal cell carcinoma or glioma (Jikuya et al., 2020; Şenbabaoğlu et al., 2016; Vuong et al., 2019; Yeo et al., 2022; Zhang et al., 2021). Prediction of outcome is restricted in CLL to cases with unmutated *IGHV* gene locus, which show significantly higher Galectin-9 expression compared to cases with mutated *IGHV* and are generally considered as the worse prognostic group of patients (Damle et al., 1999; Hamblin et al., 1999). As the Eµ-TCL1 mouse model mimics CLL with unmutated immunoglobulins (Yan et al., 2006), our data suggests treatment efficacy of anti-Galectin-9 specifically for this more aggressive form of CLL.

Our study focused on an in-depth characterization of the T cell landscape in CLL and we excluded myeloid cells from the analysis, which represents a limitation of our analysis. Immunosuppressive interactions of CLL-associated myeloid cells and T cells contribute to immune escape in CLL (Hanna et al., 2016; Jitschin et al., 2014), but their analysis requires fresh tissue samples to prevent a biased loss of myeloid cell subsets by freezing and thawing, and such samples are rarely available. As suppression of anti-tumor immunity is complex and multifactorial, obtaining a complete picture of all cell types and their interactions with the cancer cells will be necessary for the development of combinatorial treatment targeting multiple immune cell types and checkpoints to avoid immune escape and resistance to therapy. The data provided here suggest that Galectin-9 is an attractive novel therapy target for immunotherapy in CLL and likely beyond in other cancers.

## Methods

### Patient samples

Lymph node (LN, n = 21), peripheral blood (PB, n = 7), and bone marrow (BM, n = 3) samples from CLL patients, and reactive lymph node (rLN, n = 13) samples from heathy controls were obtained after informed consent and according to the guidelines of the Hospital Clínic Ethics Committee, the Ethics Committee of the University of Heidelberg, and the Declaration of Helsinki. Patients with CLL were diagnosed following the World Health Organization (WHO) classification criteria. All clinical information of the patients analyzed in this work is provided in Suppl. Table 1.

### Mice and tumor models

C57BL/6 wild-type (WT) mice were purchased from Charles River Laboratories (Germany) or Janvier Labs (France). Adoptive transfer of TCL1 leukemia cells was performed as previously described (Hanna et al., 2016; Wierz et al., 2018). In short, malignant B cells were enriched from splenocytes of Eµ-TCL1 mice using EasySep Mouse Pan-B Cell Isolation Kit (StemCell Technologies, Inc., Cologne, Germany). The CD5^+^ CD19^+^ content of purified cells was typically above 95%, as measured by flow cytometry. 1 x 10^7^ malignant TCL1 splenocytes were transplanted by intravenous (i.v) injection into 8-12 weeks-old C57BL/6 WT female mice. Leukemia progression was monitored weekly by assessment of the percentage of CD5^+^CD19^+^ cells in the PB.

### Antibody treatments

For Galectin-9 blockade experiments, mice were first transplanted with TCL1 leukemia cells. After 2 weeks, mice were assigned to different treatment arms according to the percentage of CD5^+^CD19^+^ (CLL) cells out of CD45^+^ cells in PB. Subsequently, mice were injected i.p. with 6 mg/kg of anti-Galectin-9 (clone: RG9-1) or rat IgG2b isotype control antibody (clone: LTF-2), 3 times per week for another 7 weeks. All antibodies for in vivo experiments were acquired from BioXcell (West Lebanon, NH).

### Tissue sample collection and preparation of cell suspensions

Mice were euthanized by increasing concentrations of carbon dioxide (CO_2_) or by cervical dislocation before reaching the established humane end-point. PB was drawn from the submandibular vein or via cardiac puncture and collected in ethylenediaminetetraacetic acid (EDTA)-coated tubes (Sarstedt, Numbrecht, Germany). Single-cell suspensions from spleens, bone marrow (BM) and inguinal LNs were prepared as previously described (Hanna et al., 2016; McClanahan et al., 2015). BM cells were flushed from femurs with 5 mL of phosphate-buffered saline (PBS)/5% fetal calf serum (FCS). Spleen single-cell suspensions were generated by using the gentleMACS tissue dissociator with Gentle MACS tubes C (Miltenyi Biotec). Single cell suspensions from LNs were prepared by grinding the tissue through 70 mm cell strainers (BD Biosciences, Heidelberg, Germany). Erythrocytes were lysed by using Red blood cell lysis buffer (Mitenyi Biotec).

### Flow cytometry

Single-cell suspensions were immunostained with antibodies against cell surface proteins (Key Resource Table) in FACS buffer containing 0.1 % fixable viability dye (Thermo Fisher Scientific, 65-0866-14) for 30 min at 4 _°_C. Cells were subsequently washed twice in FACS buffer, fixed with IC fixation buffer (Thermo Fisher Scientific, 00-8222-49) for 30 min at room temperature, washed twice with FACS buffer, and stored at 4 _°_C in dark conditions until being analyzed.

For transcription factor or intracellular cytokine staining, cell surface-stained cells were fixed for 30 min at room temperature with Foxp3 fixation/permeabilization buffer (Thermo Fisher Scientific, 00-5523-00 or Miltenyi Biotec, 130-093-142). After a washing step with FACS buffer, cells were permeabilized for 30 min at room temperature with 1X permeabilization buffer (Thermo Fisher Scientific, 00-8333-56). Intracellular staining with antibodies against transcription factors or cytokines was performed in 1X permeabilization buffer for 30 min at room temperature. Excess antibodies were washed twice with 1X permeabilization buffer and cells were resuspended in 1X permeabilization buffer and stored at 4_°_C in dark conditions until they were analyzed by flow cytometry.

For intracellular CD107a assessment, splenocytes were stimulated overnight with phorbol myristate acetate (PMA) and ionomycin (100 nmol/L and 1 μmol/L) and incubated for 4 hours with Brefeldin A (BFA, 1X) prior to washing and cell surface staining.

Cell fluorescence was assessed using a BD LSRFortessa^TM^ (BD Biosciences) or a Novocyte Quanteon (Agilent) flow cytometer, and data was analyzed using FlowJo X 10.0.7 software (FlowJo). In each experiment, single fluorochrome stainings were used to compensate for spectral overlap. Fluorescence minus one (FMO) controls were employed for proper gating of positive cell populations. FMO-normalized Mean Fluorescence Intensity (nMFI) for unimodal distributions or percentage of positive cells for bimodal distributions were determined for populations of interest. The standard gating strategy for analysis of CLL cells and T cells is depicted in Suppl. Figure 4A.

### Immunofluorescence of whole lymph node sections

Paraffin-embedded tissue sections of lymph nodes were obtained from the National Center for Tumor Disease (NCT), Hospital Clínic de Barcelona, and the University of Würzburg.

After validation of primary and secondary HRP-conjugated antibodies using DAB chromogenic detection, incubation times and appropriate antibody concentrations were optimized using the Opal™ 5-color kit (Akoya Biosciences, NEL840001KT). Epitope stability was assessed to determine the order of each antibody in sequence, and antigen stripping efficiency was confirmed in all used primary antibodies.

Deparaffinization and rehydration of lymph node sections was performed by immersion in xylene for 10 minutes twice, followed by immersion in a series of descending ethanol concentrations prior to distilled water. Slides were then cooked in a steam cooker with antigen retrieval buffer pH9 (Akoya Biosciences, AR900250ML) at 100 _°_C for 25 minutes. Slides were allowed to cool at room temperature and washed in tris-buffered saline containing 0.1 % tween 20 (TBS-T). Next, slides were incubated with blocking buffer (Akoya Biosciences, NEL840001KT) for 10 minutes at room temperature and incubated with the primary antibody for the indicated time and concentration (Key Resource Table). After three washing steps with TBS-T, slides were incubated with anti-rabbit HRP-conjugated secondary antibody (Akoya Biosciences, NEL840001KT). Sections underwent rounds of antigen stripping via incubation at 100 °C in AR9 buffer (Akoya Biosciences, AR900250ML) for 25 minutes, allowing removal of the excess of antibody complex formed. For multi-color staining, tissue sections went through multiple rounds of blocking, primary and secondary antibody labelling, each round with a different Opal^TM^ dye (Akoya Biosciences, NEL840001KT). Finally, background signal reduction through incubation for 10 minutes at RT in 0.1 % Sudan Black (Sigma-Aldrich, 199664) took place and sections were washed thoroughly in TBS-T. Finally, nuclei were stained with DAPI (Akoya Biosciences, NEL840001KT) and slides were mounted using ProLong Diamond Antifade Mountant (Life Technologies, P36965).

Slides were imaged using a Carl Zeiss Axioscan 7 slide scanner at 20X.

### Immunofluorescence staining and imaging of CLL TMA

TMA 159/2 was de-paraffinized and subjected to antigen retrieval for 5 minutes at 95°C followed by 5 minutes at 107°C, using pH8.5 EZ-AR 2 Elegance buffer (BioGenex). To reduce tissue autofluorescence, slides were placed in a transparent reservoir containing 4.5% H_2_O_2_ and 24 mM NaOH in PBS and illuminated with white light for 60 minutes followed by 365 nm light for 30 minutes at room temperature as previously described (Lin et al., 2018). Slides were rinsed with surfactant wash buffer (0.025% Triton X-100 in PBS), placed in a humidified stain tray, and incubated in Image-iT™ FX Signal Enhancer (Thermo Fisher) for 15 minutes at room temperature. After rinsing with surfactant wash buffer, the slides were placed in a humidity tray and stained with the panel of fluor- and hapten-labeled primary antibodies in PBS-Antibody Stabilizer (CANDOR Bioscience GmbH) containing 5% mouse serum and 5% rabbit serum for 2 hours at room temperature. Slides were then rinsed again with surfactant wash buffer and placed in a humidified stain tray and incubated with SYTOX Blue (Thermo Fisher), ArgoFluor™ 845 mouse-anti-DIG in PBS-Antibody Stabilizer containing 10% goat serum for 30 minutes at room temperature. The slides were then rinsed a final time with surfactant wash buffer and PBS, coverslipped with ArgoFluor™ Mounting Media (RareCyte, Inc.) and dried overnight (Lin et al., 2022).

Slides were imaged using an Orion instrument (RareCyte, Inc.) at 20X. Raw image files were processed to correct for system aberrations; then signals from individual targets were isolated to separate channels using the Spectral Matrix obtained with control samples, followed by stitching of FOVs to generate a continuous open microscopy environment (OME) pyramid TIFF image.

### Image analysis

#### Preprocessing

TMA 159/2 with 18-color staining was analyzed using the multiple-choice microscopy pipeline MCMICRO (Schapiro et al., 2022) (Commit 91fcd5eefd69da112414b06cf3a65a9a66afeccf). Dearraying was performed by Coreograph (Schapiro et al., 2022) 2.2.8, Unmicst (Yapp et al., 2022) 2.7.0 segmented the cells based on the nuclear channel Sytox, S3segmenter (Saka et al., 2019) 1.3.12 generated single-cell masks and Mcquant (Schapiro et al., 2022) 1.5.1 quantified the signal of each of the 18 channels in each cell. The corresponding parameters (params.yml) for full reproducibility of this MCMICRO runs are available on Github (https://github.com/SchapiroLabor/ImageAnalysisTMA159_2).

Following visual inspection of Coreograph’s QC file, all cores that were damaged were filtered out. Additionally, the 4 muscle tissue cores – which were used as control – were filtered. In total we performed the downstream analysis with 42 cores. Channel numbers 2 (Sytox), 14 (E-cadherin), 17 (Pan-CK) and 18 (AF-Tissue) were removed for downstream analysis since (i) Sytox and AF-Tissue represent nuclei and autofluorescence which are not relevant for cell type calling; and (ii) E-cadherin and Pan-CK expression are not expected in LN tissue and therefore, failed our visual QC due to unspecific staining patterns. Image visualization was performed with QuPath (Bankhead et al., 2017) and Napari (Chiu et al., 2022).

#### Batch correction

Cell type clustering on the raw data results in clusters consisting mainly of cells of a single core of origin. This is due to preanalytical variability, which requires batch correction on core-level. Therefore, marker intensities were normalized with Mxnorm (Harris et al., 2022) 0.0.0.9000 (transform: log10_mean_divide, method: none) before using Rphenograph (Levine et al., 2015) 0.99.1 to cluster the individual cell types.

#### Cell type assignment

Cell types were assigned based on marker thresholding (positive / negative) and comparison to an expert curated list of markers associated with the specific cell types (Suppl. Table 2).

#### Spatial Analysis

Neighborhood analysis was performed with Giotto 1.1.2 (Dries et al., 2021). After building a Delaunay network and running 1000 simulations, we compared the number of observed interactions between the cell types with their respective expected number of interactions. Cell-cell interactions between the same cell types are significantly enriched because of segmentation inaccuracies. Cells are slightly over segmented creating multiple cells with the same phenotype.

#### Statistical analysis

The subsequently calculated cell type frequencies in each core were used to perform two-sided t-tests (base package stats) for changes in cell type frequencies between patients that were positive or negative for a variety of binary clinical parameters (sex, IGHV, del13q, del17p, died, treated, del17p_tp53 (del17p and/or tp53 positive)). Additionally, Cox regression (survivalAnalysis 0.3.0) was calculated on overall survival and time to next treatment. Finally, correlation with the Pearson method was checked (cor.test() in base package stats) between overall survival, time to next treatment and all cell type frequencies. Up to the statistical analysis, calculations were performed under R version 4.1.0 (2021-05-18) on platform x86_64-apple-darwin17.0 (64-bit) running under macOS 13.0. Statistical analysis was done on R version 4.0.5 (2021-03-31) on platform: x86_64-apple-darwin17.0 (64-bit) running under: macOS Big Sur 10.16.

All mentioned steps of image processing were performed on the BWforCluster.

### Mass cytometry

A panel of 42 heavy metal-labeled antibodies for detection of both surface and intracellular proteins was adopted from Bengsch et al. and modified based on literature research in order to characterize diverse T cell phenotypes (Bengsch et al., 2018). The complete list of proteins detected and the heavy metal-conjugated antibodies used are listed in the Key Resource Table. For most of the markers, heavy metal-conjugated antibodies were commercially available and purchased from Fluidigm. Where no heavy metal-conjugated antibodies were commercially available, coupling of heavy metal to respective antibodies was performed in-house using the Maxpar® X8 Multimetal Labeling kit (Fluidigm) following the manufacturer’s instructions. ^113^In, ^115^In and ^139^La heavy metal isotopes (#) were not available from Fluidigm and were purchased from Trace Sciences International (^113^In) and Sigma (^115^In, #203440-1G; ^139^La, #211605-100G) Following the preparation of a 1M solution, ^113^In, ^115^Ln and ^139^La heavy-metal isotopes (#) were conjugated to monoclonal purified IgG antibodies using the Maxpar® X8 Multimetal Labeling kit (Fluidigm, #201300).

Frozen cell suspensions were thawed and washed prior to incubation in 15 mL of pre-heated RPMI 1640 containing 10 % FBS in a roller incubator for 30 min at room temperature. Cells were filtered to remove dead cells and B-cell depletion was performed using human CD19 microbead (Miltenyi Biotec) labeling according to the manufacturer’s protocol. Unlabeled CD19^-^ cells were collected, washed, filtered, and counted prior to mass cytometry processing.

CD19-depleted single-cell suspensions were stained as previously described (Wierz et al., 2018). Briefly, cells were stained with 5 μM cisplatin (Cell-ID Cisplatin, Fluidigm) for 5 min in order to label dead cells. Cells were then washed once and cell surface staining was performed for 30 min at room temperature. After a washing step, cells were fixed with Fixation/Permeabilization buffer (Thermo Fisher Scientific) following the manufacturer’s instructions. Intracellular staining was performed by incubating the antibody-cocktail for 30 min at room temperature. After a washing step, cells were stained with cell-ID Intercalator-Ir (Fluidigm) in fixation and permeabilization solution, followed by another two washing steps with PBS and ddH_2_O, respectively. Prior to acquisition, cells were resuspended at a concentration of 5 x 10^5^ cells/mL in ddH_2_O with 1:10 calibration beads (EQ Four Element Calibration Beads, Fluidigm). Samples were analyzed at a flow rate of 0.03 mL/min with the Helios mass cytometer (Fluidigm) of the National Cytometry Platform (LIH). Initial data processing and quality control were performed. Flow cytometry standard (FCS) files were normalized with EQ beads using the HELIOS instrument acquisition software (Fluidigm).

As a first analysis step, samples were pre-gated as follows: gates were placed on cells (Beads vs. Ir191), singlets (Ir193 vs. Ir191) and live cells (Pt195^-^). Next, non-B immune cells (CD19^-^ CD45^+^) and CD3^+^ T cells (CD3^+^ NCAM^-^) were selected exported as .fcs files.

Raw data .fcs files were merged into a flowSet object using flowCore package (Hahne et al., 2009). Signal intensities for each marker were arcsinh-transformed using a co-factor of 5 (default) (Bruggner et al., 2014). Sample quality was assessed based on cell counts per sample and samples with less than 1,500 cells were excluded. Multi-dimension scaling (MDS) plotting using median arcsinh-transformed marker expression for all cells in each sample was used for sample clustering analysis.

Cytometry data analysis tools (CATALYST) R package (Chevrier et al., 2018) containing FlowSOM (Van Gassen et al., 2015) and ConsensusClusterPlus (Wilkerson and Hayes, 2010) metaclustering methods was used for cell cluster identification using all cells from all samples. Clustering was performed using arcsinh-transformed expression of 33 markers, i.e., excluding the markers DNA content and cisplatin viability, which were used to select viable, single cells, and CD19 and HLA-DR, which were used to exclude B cells (Key Resource Table). TIM3, LAG3, GITR, CD47 and 4-1BB were excluded from the analysis due to low signal and likely adding noise in the cluster generation process.

The maximum number of clusters allowed to be evaluated was set to maxK = 30, after biological relevance of obtained clusters was assessed, and k < 30 and k > 30 were verified to be underfitting or overfitting, respectively. Cell clustering was visualized using the UMAP algorithm, displaying 1 x10^3^ random cells from each sample. The robustness of this approach was evaluated by repeatedly plotting an increasing number of randomly picked cells from each sample, with a range of 200 to 1,000 cells, and evaluating the similarity of the 30 subpopulations recognized in the respective UMAP plots. Sample #HD3 was excluded as an outlier.

### Cell sorting for single-cell RNA and TCR sequencing

Single-cell suspensions from LNs of CLL patients were retrieved by partially thawing vials of cryopreserved cells in order to preserve unused cells. Samples were stained for cell surface proteins for 30 min. After washing with PBS, cells were resuspended in PBS + 5 % FBS containing 0.2 μg/mL DAPI prior cell sorting. The gating strategy for CD3^+^ T cell and CLL cell sorting is depicted in Suppl. Figure 5A. Cells were sorted in PBS + 2 % FBS using a BD FACSAria^TM^ II or BD FACSArisa Fusion (both from BD Biosciences) cell sorters. The purity of cells after sorting was above 95 %.

### Single-cell RNA sequencing library construction using the 10x Genomics Chromium platform

Five CLL LN samples were processed for single-cell TCR and 5’ gene expression profiling of CD3 T cells and CLL cells using the Chromium Next GEM Single Cell V(D)J Solution from 10x Genomics following the manufacturer’s instructions. Briefly, for a target cell recovery of 5,000 cells, the concentration of T cells was adjusted to 1 x10^3^ cells/μL and tumor cells were added at a ratio of 1:20. Cells were then loaded into the Single Cell A Chip (10x Genomics). GEX libraries were sequenced on a HiSeq 4000 machine (Illumina) or on a NovaSeq 6000 (Illumina), and V(D)J-enriched libraries were sequenced on a NextSeq 550 (Illumina).

### Single cell RNA-Seq data processing

Sequencing reads were aligned by Cell Ranger (5.0.1) to reference version hg38(2020-A) for human data and to reference version mm10(2020-A) for mouse data. Raw count matrices were processed by the quality control pipeline to determine the optimal cut-offs for filtering. Cells were filtered by read counts and mitochondrial RNA content. Lower cut-off for read counts were determined by read-counts distribution for samples in each dataset and were set at 350 for human and 500 for mouse, mitochondrial RNA content should be lower than 10%.

Raw gene count matrices were processed by Seurat(v4.1.0) (Hao et al., 2021), expression values were normalized and scaled by SCTransform. Doublets were filtered by DoubletFinder(2.0.3) (McGinnis et al., 2019). Dimensional reduction was performed by Principle Component Analysis (PCA). Datasets were integrated by HARMONY (Korsunsky et al., 2019) to reduce batch bias on calculated principal components. Sample identity was used as the batch variable for HARMONY integration. Uniform Manifold Approximation and Projection (UMAP) was used for dimensional reduction on first 23 HARMONY computed cell-embeddings.

Cluster were defined by the Louvain method at the resolution of 0.5, clusters defined at this step were used for annotation by manual curation. Markers for each cluster were produced by differential expression analysis searching for overexpressing markers of each cluster. Top 10 overexpressing markers of each cluster were illustrated by a heatmap. To reveal potential substructure among cell types of interest, we performed a subcluster analysis on the T_FH_ cluster. UMAP analysis was repeated on the harmony corrected embeddings of cells from the T_FH_ cluster to reveal substructures.

### Pseudotime analysis

Diffusion pseudotime analysis was performed using destiny(3.8.1) (Angerer et al., 2016). Cells expressing CD4 and CD8 were analysed separately. To retrieve the pseudotime of each marker differentiated cell groups, “DPT” function from destiny was used for computing the diffusion pseudotime on each cell. For each cell type, the root nodes were placed within the corresponding naïve cell type cluster.

To performed differential gene expression analysis on pseudotime, the diffusion pseudotime values of cells in trajectories were analysed together with their transcript expression values by tradeSeq (Van den Berge et al., 2020). Differentially expressed genes were plotted against ranked diffusion pseudotime for illustration.

### TCR data analysis

V(D)J transcripts from single cells were aligned and counted using the Cell Ranger pipeline (5.0.1). GRCm38 VDJ Reference 5.0.0 from 10X genomics was used for mouse data and GRCh38 VDJ Reference 5.0.0 from 10X genomics was used for human data. In addition, an output file containing TCR α- and β-chain CDR3 nucleotide sequences and a cell barcode for all single cells was generated. Only productive rearrangements were evaluated, and 3 or more cells containing the same α- and β-chain CDR3 consensus nucleotide sequences were considered cell clones. scRepertoire (Borcherding et al., 2020) was used to process contig and clonotype information from Cell Ranger. The summarized TCR information per cell was merged with Seurat object using the shared cell barcodes between the TCR library and the RNA-Seq library for integrated analysis. Shannon diversity index was computed to represent the diversity of the TCR repertoire in each single cell clusters. Cells with expanded TCRs are categorized by their expansion level and then illustrated by umap plots. To illustrate expanded TCRs shared between different clusters, cells with expanded TCRs were represented by two types of chord diagrams. The first type depicts how other clusters share expanded TCRs with CD8 TEM clusters, where a chord diagram was created for each CD8 TEM cluster. Circular fragments in these chord diagrams represent cells carrying an expanded TCR in each cell type, arcs connect cells that share the same TCRs with the CD8 TEM of interest. Width of the arcs is scaled by the number of cells sharing same TCRs. The second type of chord diagram present a similar concept, it depicts TCR sharing across all cell types instead of one.

The VDJdb browser (Bagaev et al., 2020) was used in order to identify CDR3 regions from the V(D)J single-cell dataset whose antigen-specificity has been reported. Both TCR α- and β-chain CDR3 amino acid sequences from each sample were used for the literature analysis aimed at identifying previously described binding epitopes.

### Ligand-receptor interaction analysis

To find differential ligand-receptor interactions between datasets, a control dataset derived from 5 reactive lymph nodes (Aoki et al., 2020) was added for comparison. The reactive lymph node dataset was independently processed by the same quality control pipeline and single cell analysis pipeline. CellChat was used for differential cell-cell communication analysis. Ligand-receptor interactions from OmnipathR database (Türei et al., 2021) were included as candidates. CellChat analysis was first performed independently on the two datasets using LIANA (Dimitrov et al., 2022). The results were then contrasted by CellChat for differential interactions. Labels from the two comparing datasets were harmonized to 8 major cell types (CLL/B, CD4 T_CM_, T_REG_, CD4 T_EM_, CD8 T_EM_, CD4 T_N_, CD8 T_N_, T_FH_) for comparison. Differential Interactions with p-values below 0.01 were visualized by the dot plot functions of CellChat.

### Expression of Galectin-9 protein and its impact on survival in CLL

Galectin-9 protein levels in CLL patient samples were obtained from Herbst SA, Vesterlund M, *et al*. (Herbst et al., 2022). Mean relative Galectin-9 levels were compared between IGHV-mutated and -unmutated CLL patients using a two-sided Wilcoxon signed-rank test. Time to next treatment (TTNT) was calculated from the date of sample collection to subsequent treatment initiation. Proportional hazards regression (Cox regression) was used to calculate the impact of Galectin-9 abundance on TTNT using the R package survival (version 3.2-3). Kaplan-Meier curves were plotted for all samples and for IGHV-mutated and -unmutated CLL patients separately.

### Expression of LGALS9 and its impact on survival in TCGA samples

Expression of *LGALS9* in TCGA samples was gathered with R (4.3.1) and Bioconductor using library RTCGA.rnaseq (1.3.0,2015-11-01) and RTCGA.clinical (1.3.0,2015-11-01). Survival curves were prepared by survminer (0.4.9) splitting the samples based on the median *LGALS9* expression into two groups. Differences in survival were assessed using Cox proportional hazard model and multiple testing was corrected according to Benjamini Hochberg. GEPIA2 was used to compare tumor and normal samples from TCGA and the GTEx projects, using a standard processing pipeline (Tang et al., 2019).

### Expression of LGALS9 in single-cell RNAseq data sets

The expression of *LGALS9* was assessed in renal cell carcinoma (SCP1288) (Bi et al., 2021) and glioblastoma (SCP1985) (Abdelfattah et al., 2022). Data was collected from (https://singlecell.broadinstitute.org/single_cell) and processed in R using Seurat (4.9.9.9044). In short, data was loaded using Read10X and after the creation of a Seurat object (min.cells=3, min.features=200) processed using default values with NormalizeData, FindVariableFeatures, ScaleData, RunPCA, FindNeighbours, FindClusters, and RunTSNE. Figures were prepared using FeaturePlot and DimPlot highlighting cell types. Violin plots were prepared using ggplot2 (3.4.2).

## Data availability

Human single-cell RNA- and TCR-sequencing data are deposited to European Genome-phenome Archive (EGA) under the accession ID (Pending) and are available on request. Mouse single-cell RNA- and TCR-sequencing data are available on Gene Expression Omnibus under the accession ID (Pending).

## Acknowledgements

We would like to thank all members of the High Throughput Sequencing Unit of the Genomics and Proteomics Core Facility, of the Light Microscopy Core Facility, and of the Single-cell Open Lab (scOpenLab) at the German Cancer Research Center (DKFZ, Heidelberg) for their technical assistance. We also thank the National Cytometry Platform (NCP) of the Luxembourg Institute of Health (LIH, Luxembourg) for assistance with the generation of cytometry data, in particular Maira Konstantinou and Dominique Revets. The NCP is supported by funding from Luxembourg’s Ministry of Higher Education and Research (MESR). Tissue microarrays and lymph node sections were provided by the Tissue Bank of the National Center for Tumor Diseases (NCT) Heidelberg, Germany in accordance with the regulations of the tissue bank and the approval of the ethics committee of Heidelberg University.

This study was supported by a research grant of the German Cancer Aid (Deutsche Krebshilfe) to MS, by grants from FNRS-Télévie to IFB (7.4529.19, 7.6603.21), MW (7.6504.18), GP (7.4501.18, 7.6518.20), SG (7.4502.19, 7.6604.21), and from the Luxembourg National Research Fund (FNR), Fondation Cancer and Plooschter Projet to EM and JP (C20/BM/14582635, and C20/BM/14592342). The authors acknowledge support by the state of Baden-Württemberg through bwHPC and the German Research Foundation (DFG) through grant INST 35/1597-1 FUGG. MC, KB and DS are supported by the German Federal Ministry of Education and Research (BMBF 01ZZ2004).

## Author information

L.L.C. designed the study, performed experiments, analyzed data, generated figures, and wrote the paper. J.K.L.W. performed bioinformatical analyses, generated figures, and wrote the paper. I.F.B. and M.W. conceptualized part of the study, performed experiments, and analyzed data. Y.P. analyzed scRNA-seq data and CyTOF data. A.F. conducted experiments, analyzed data, and generated figures. S.G., G.P., C.S. A.C., F.C., P-M.B. and D.E.C. performed experiments and analyzed data. M.C., K.B. and D.S. analyzed multiplex imaging data, and generated figures. M.I. performed bioinformatical analyses and generated figures. T.R. and S.D. provided input regarding cluster annotations and performed correlation analyses. J.P.M. established the pipeline for single-cell RNA-sequencing. E.G.-H. and A.R. supported the study with lymph node sections and expertise in interpreting imaging data of lymph nodes. D.C., E.C., and S.D. provided clinical samples and information. P.L. provided logistic and budget support and was involved in scientific discussions. E.M., J.P., M.Z., and M.S., conceptualized the study, guided experiments and data analysis, provided logistic and budget support, and wrote the paper.

## Ethics declarations

Samples from patients and heathy controls were obtained after informed consent and according to the guidelines of the Ethics Committees of the involved University hospitals and the Declaration of Helsinki.

All experiments involving laboratory animals were conducted in pathogen-free animal facilities at the German Cancer Research Center in Heidelberg or the Luxembourg Institute of Health in Luxembourg with the approval of the Regierungspräsidium Karlsruhe (G-77/19 and G-112/21) and the Luxembourg Ministry for Agriculture (#LUPA 2019/21), respectively. Mice were treated in accordance with the European guidelines.

## Supplemental Excel Table Titles

**Suppl. Table 1:** Information on patients and donors

**Suppl. Table 2:** Cell cluster annotations

**Suppl. Table 3:** Correlations with clinical data

**Suppl. Table 4:** VDJdb results of TCR-seq data

