## Supplementary figures and images for "High-dimensional single-cell definition of CLL T cells identifies Galectin-9 as novel immunotherapy target"

### Supplementary Figure 1

#### Supplementary Figure 1

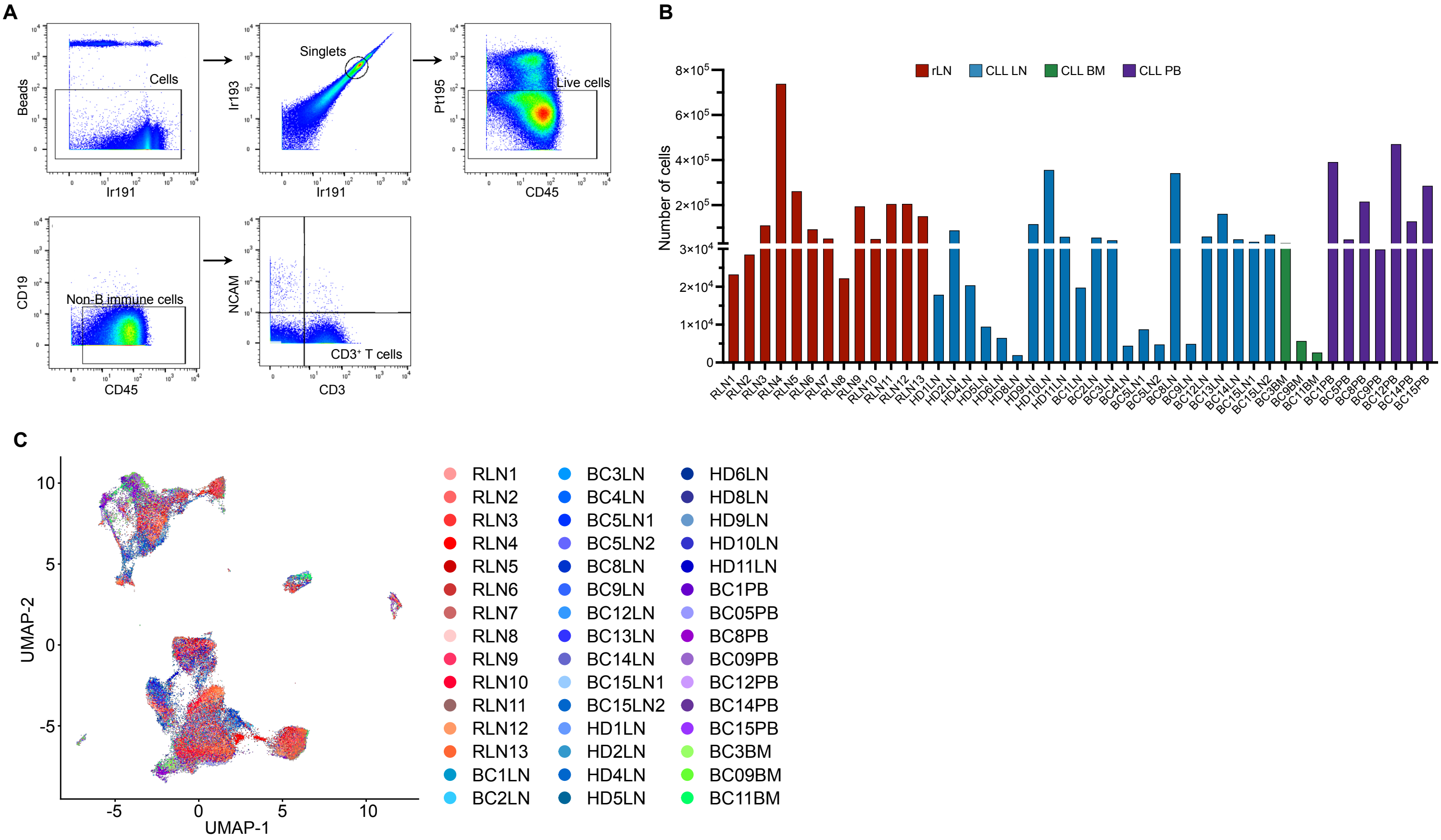

### Supplementary Figure 2

Supplementary Figure 2

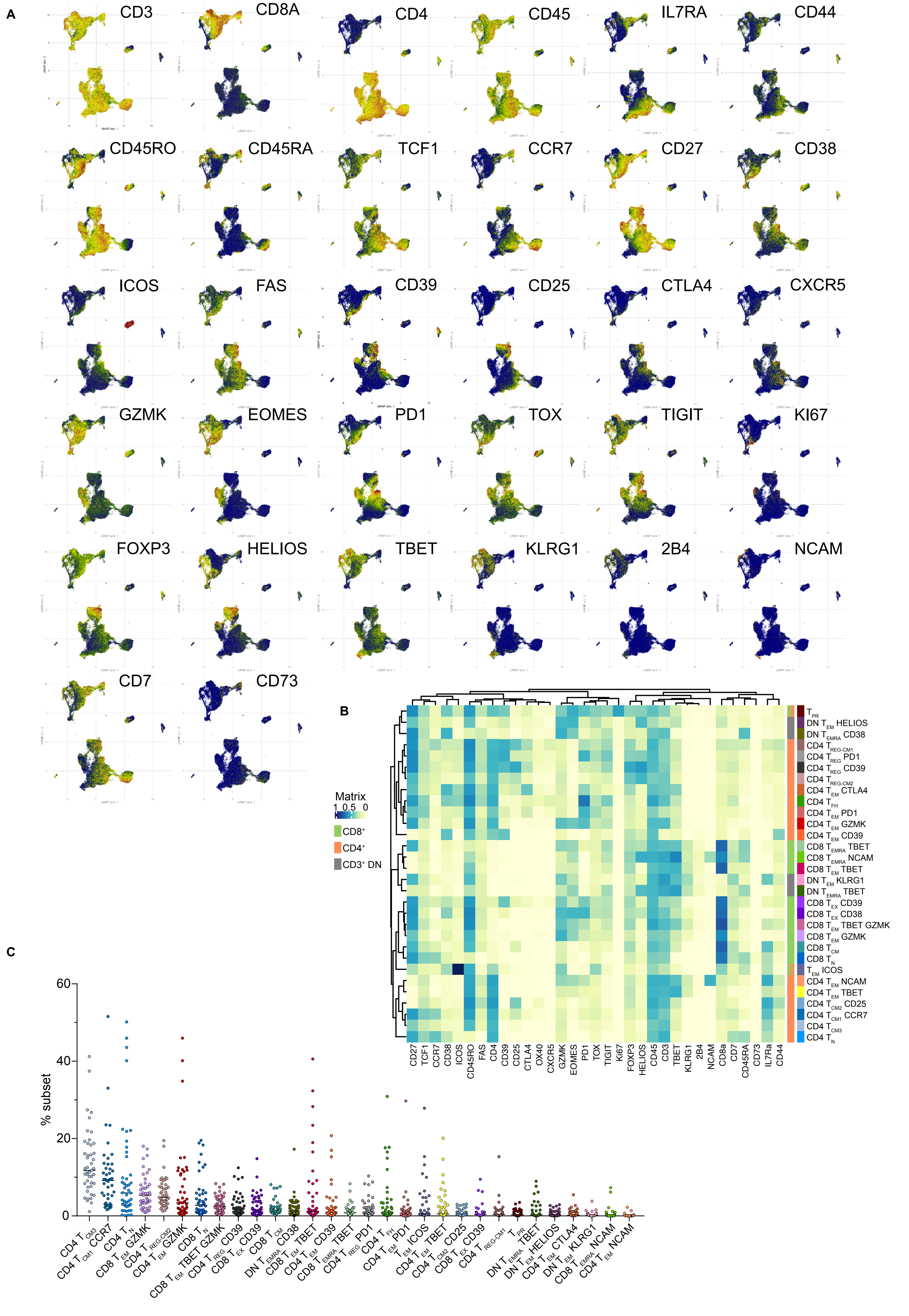

### Supplementary Figure 3

Supplementary Figure 3

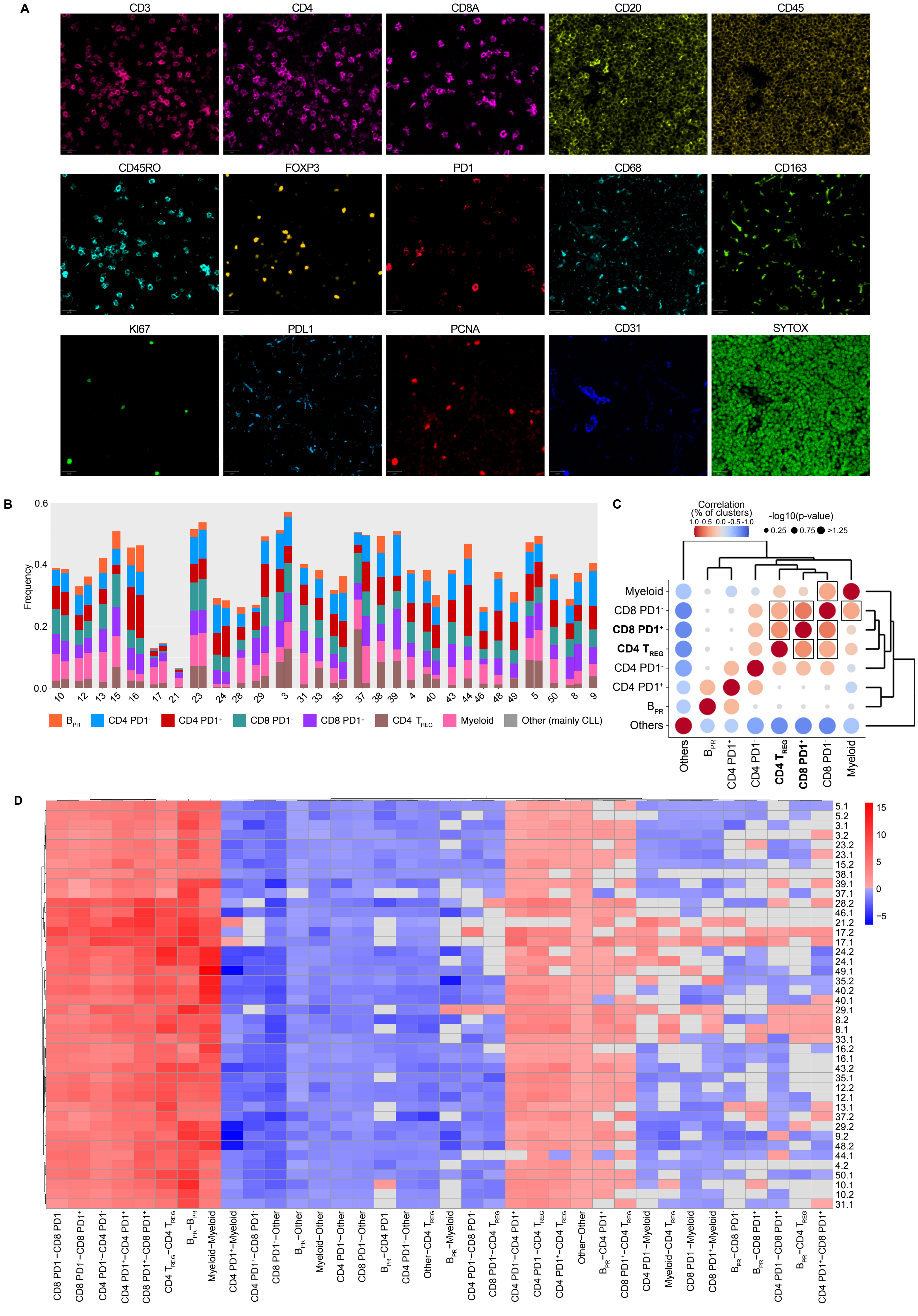

### Supplementary Figure 4

Supplementary Figure 4

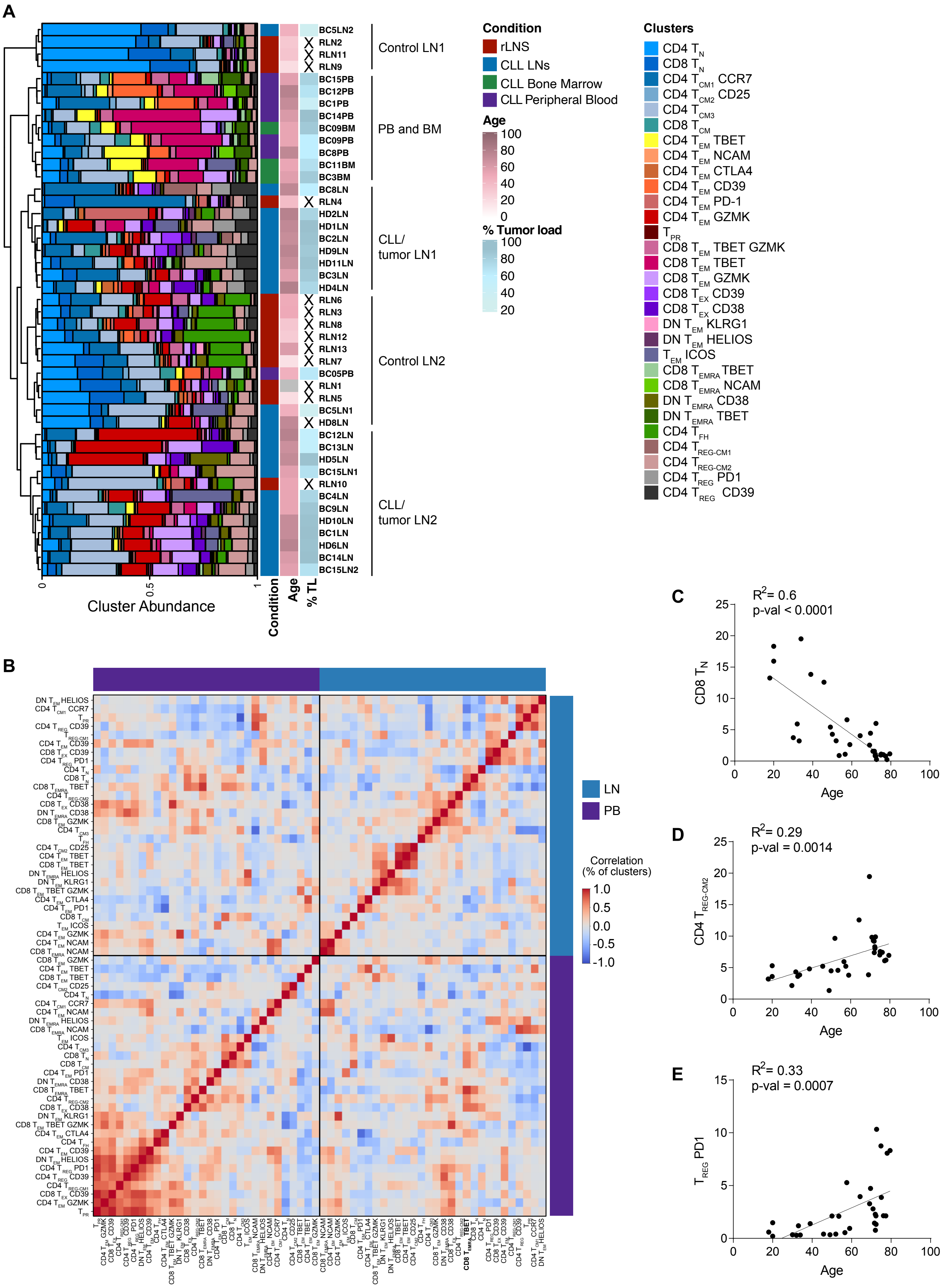

### Supplementary Figure 5

Supplementary Figure 5

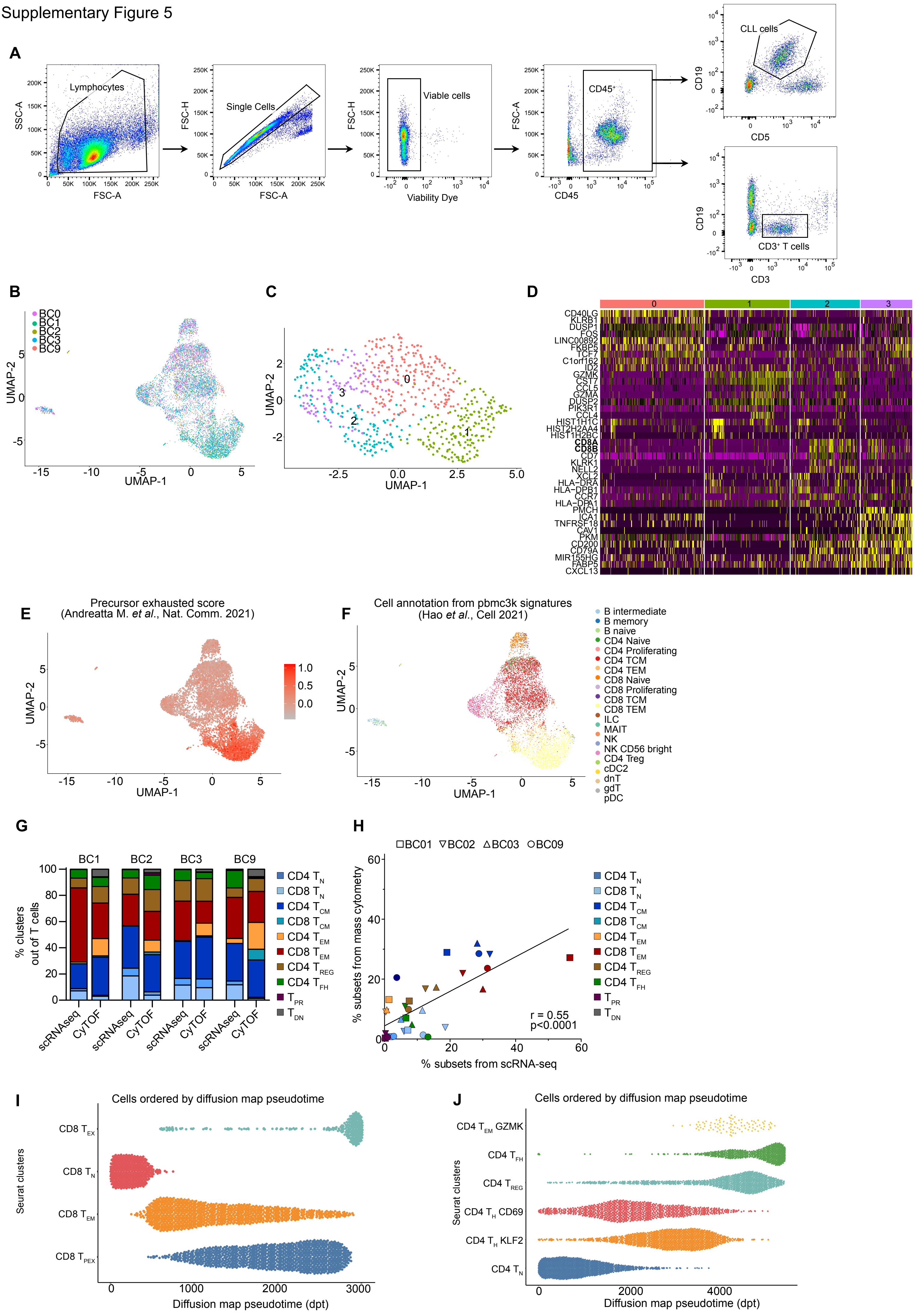

### Supplementary Figure 6

Supplementary Figure 6

**A**

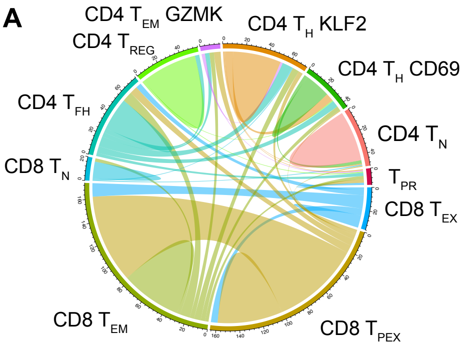

### Supplementary Figure 7

Supplementary Figure 7

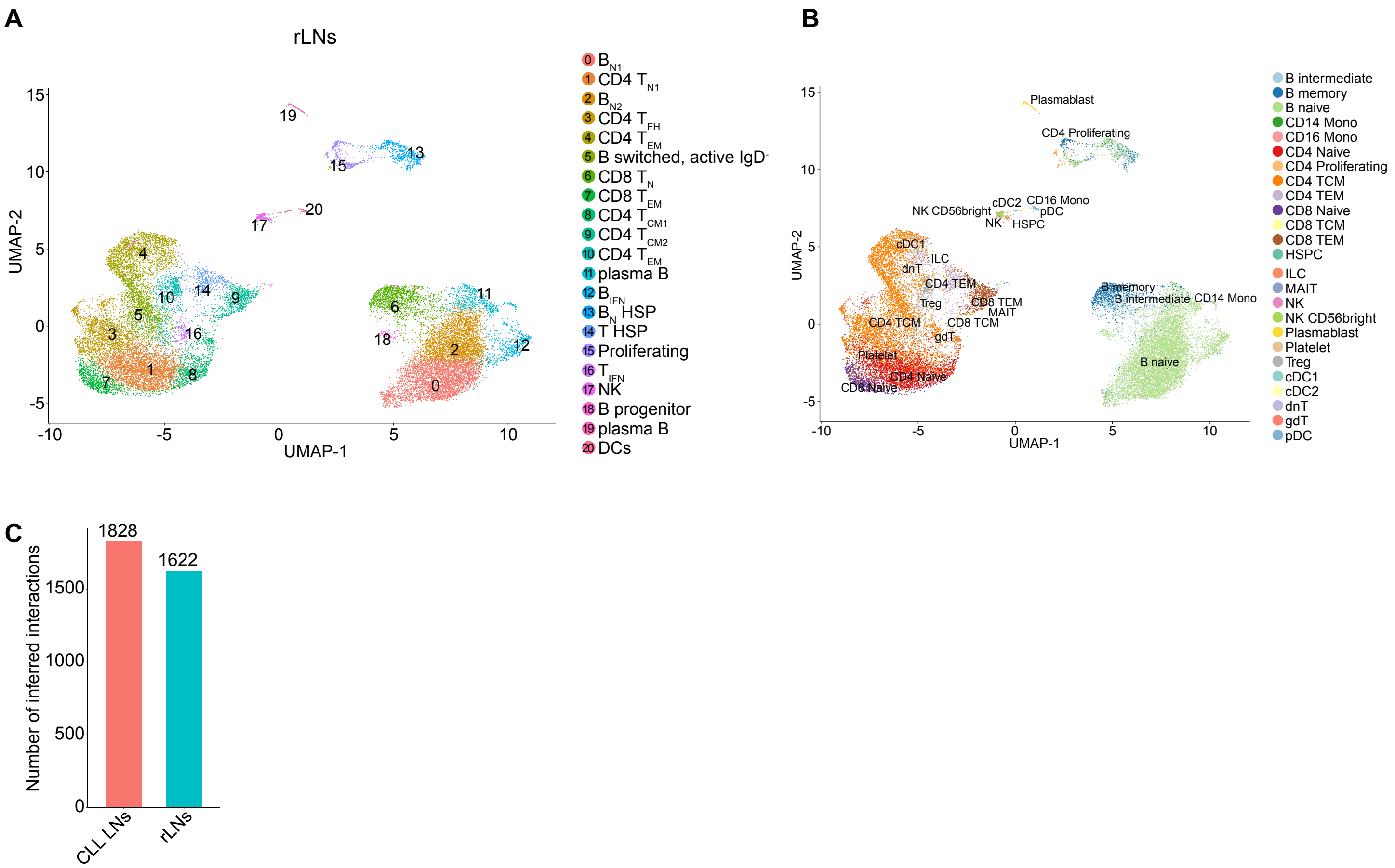

### Supplementary Figure 8

Supplementary Figure 8

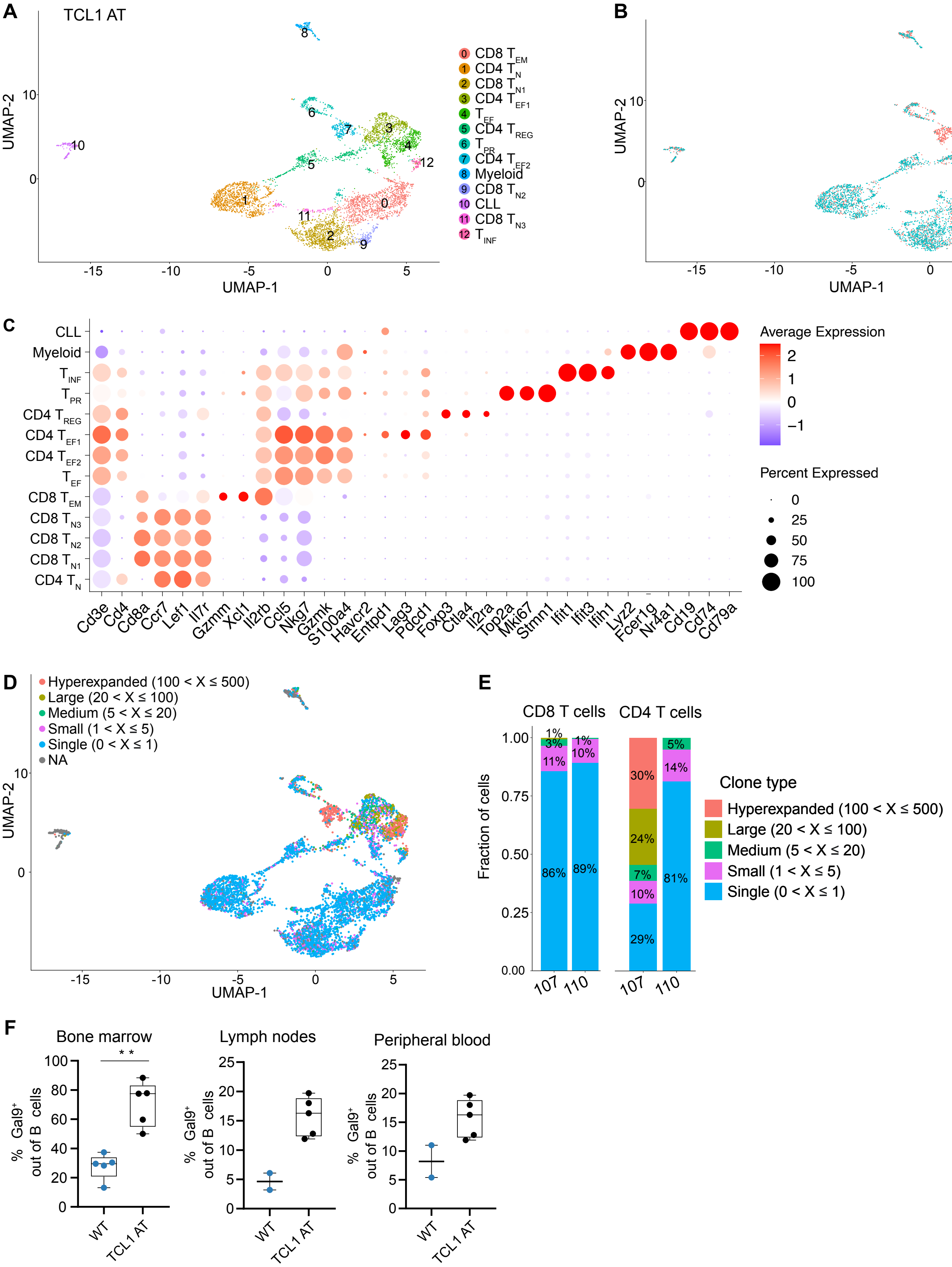

### Supplementary Figure 9

Supplementary Figure 9

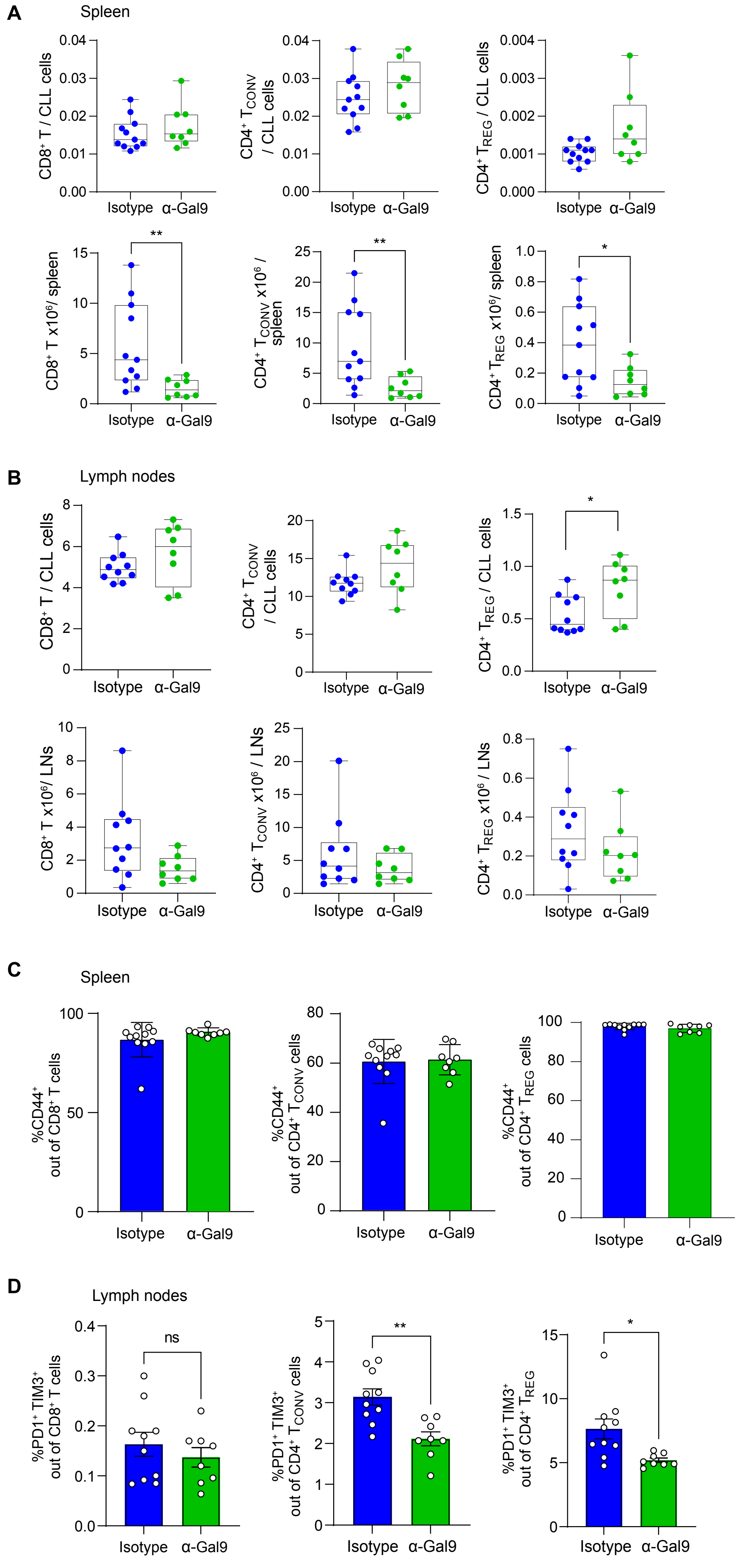

### Supplementary Figure 10

Supplementary Figure 10

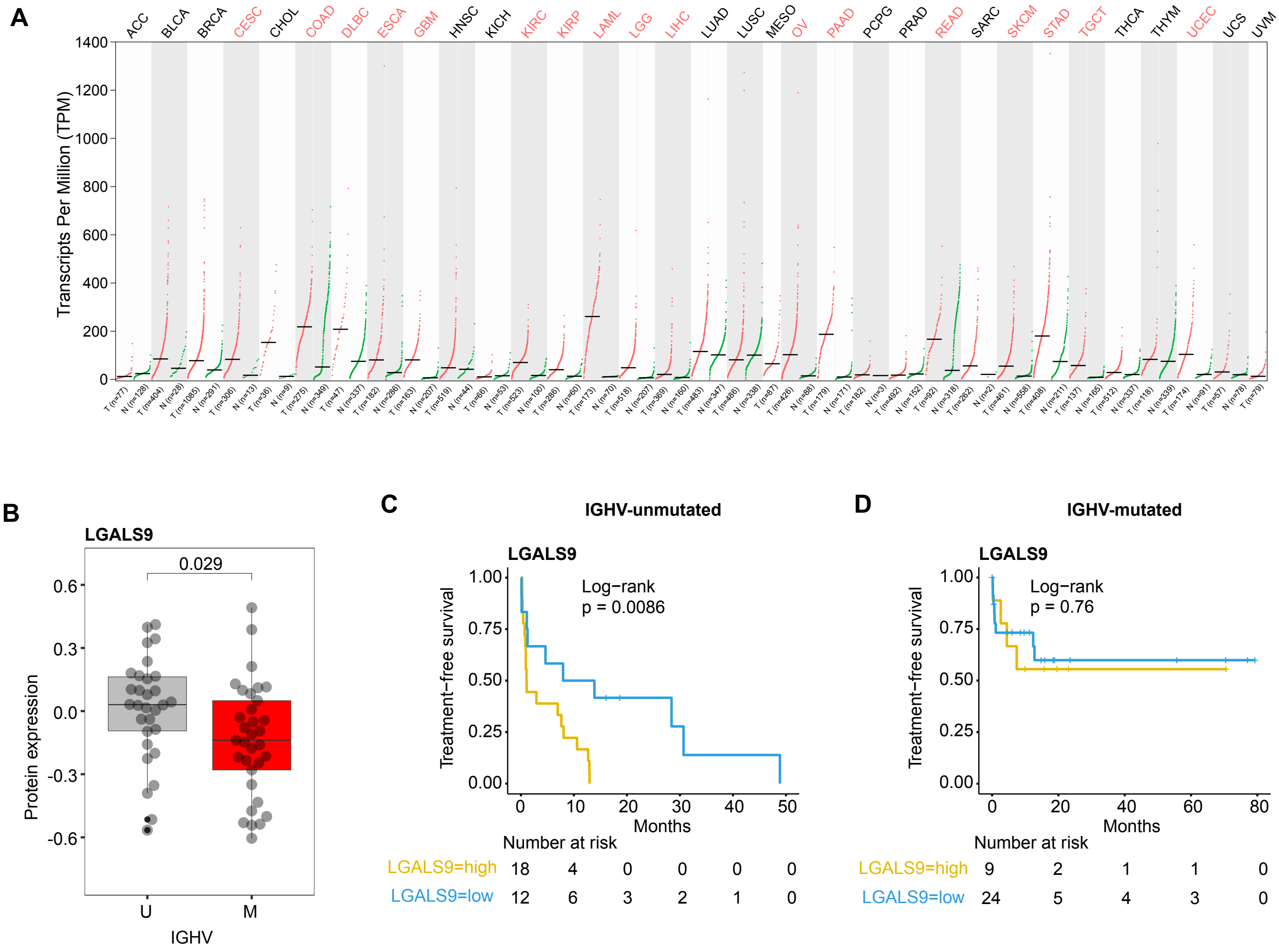
